# Psychopathic traits predict neural responses to emotional movies in the general population

**DOI:** 10.1101/2023.07.31.551268

**Authors:** Anna Aksiuto, Juha M. Lahnakoski, Janne Kauttonen, Heini Saarimäki, Sofia Volynets, Lauri Nummenmaa, Mikko Sams, Iiro P. Jääskeläinen

## Abstract

Psychopathic personality traits such as callousness, manipulativeness, and lack of empathy are continuously distributed in non-institutionalized populations. Here, we show that brain responses to emotional movies in healthy females vary as a function of primary (PP) and secondary psychopathy (SP) traits. Healthy female volunteers (n=50) with variable scorings in Levenson Self-Report Psychopathy Scale watched a set of emotional movie clips (n=39) during functional magnetic resonance imaging of brain activity. We found that participants low on primary and secondary psychopathy traits showed higher cortical and subcortical activity when watching emotional movies, what possibly indicates the somatosensory simulation and empathizing with the emotions of movie characters. On the other hand, the individuals with high primary and secondary psychopathy traits presented a lack of vicarious engagement during watching movies and they manifested brainstem-based heightened response towards thread-related stimuli, which could indicate potent autonomic reactivity. Overall, our results indicate that variability in psychopathic traits in the general population is manifested as differential neural responses to emotional signals, and confirm previous findings of there being separate PP and SP personality characteristics due to the dissociable brain activity patterns while viewing emotional movies.

## INTRODUCTION

Psychopathic personality traits such as callousness, manipulativeness, and lack of empathy are continuously distributed in clinical, forensic, and non-institutionalized samples (De Oliveira-Souza et al., 2008; Neumann & Hare, 2008) and are known to be connected to behavioral, psychophysiological and brain activation anomalies during emotional processing (Brook, Brieman & Kosson, 2013; Casey et al., 2013). Empirical evidence suggests that it is possible to identify psychopathy subtypes based on developmental background and the extent of affective deficits such as impulsivity, anxiety, and borderline-narcissistic traits (Karpman, 1941; Skeem et al., 2003). Psychopathy is associated with phenotypic variability in personality traits, symptomatology, patterns of aggression (instrumental/impulsive), degrees of temperamental fearlessness and inhibitory control (Fowles & Dindo, 2006). It has been also shown to present different outcomes associated with the diagnosis and amenability to treatment (Kosson et al., 1990; Kennealy et al., 2010). Therefore, it was previously suggested that two distinctive subgroups of psychopathy exist (Blagov et al., 2011; Markus et al., 2013).

Primary psychopathy (PP; also known as Factor 1 features) is associated with callous-unemotional traits and narcissism with grandiose and antisocial behavior. Secondary psychopathy (SP; also known as Factor 2 features) is linked with criminality, social deviance and impulsivity. It qualifies more often for borderline personality disorder and vulnerable narcissism with reactive aggression (Frick et al., 2000; Vaillancourt & Brittain, 2019). The individuals with SP have been observed as neurotic and prone to experiencing anxiety, depression, distress, and anger (Hicks & Patrick, 2006; Kimonis et al., 2011). However, in contrast to PP, SP was shown more responsive to punishment (Newman et al., 1992). Furthermore, the PP traits have been connected to restlessness and impatience and SP to problems in interest and stimulation (Wink & Donahue, 1997). Importantly, it has been suggested that PP is predominantly congenital whereas SP develops environmentally via early-life traumas and social disadvantage (Frazier, Ferreira & Gonzales, 2019).

Among the abnormalities connected to psychopathy disorder, the critical one surrounds emotional processing. Psychopathy is believed to develop from specific deficits in recognizing fear and sadness (Eisenbarth et al., 2008; Munoz, 2009). However, some research reported also abnormalities in perceiving disgust (Hansen et al., 2008), anger (Blair, 2018), happiness (Hastings et al., 2008), surprise (Fairchild et al., 2009), guilt (Johnsson et al., 2014), excitement (Bjork et al., 2012), or negative affect in general (Patrick, 1994; Blair et al.,1997). It has been indicated that PP individuals possess muted psychophysiological responses to threat and fear (Dawel et al., 2012) as well as a pattern of affective deficits with low levels of fear, guilt and anxiety combined with high levels of callousness (Verona, Patrick & Joiner, 2001). In contrast, SP individuals are able to experience social emotions such as love, guilt and empathy (Bronchain, 2020) and show stronger autonomic arousal and self-reported reactions to negative stimuli, particularly threat (Dargis & Koenigs, 2018). Further, PP in contrast to SP, is suggested to have improved ability at identifying emotional states of others that supports successful manipulation (Brook & Kosson, 2013).

Brain basis of psychopathy in females is poorly understood as the condition is more commonly reported in the male population. Nevertheless, it is suggested that the prevalence rates among female inmates are approximately half (15.5%) that of males in prison (25-30%; Salekin et al., 1997). In the general population, it is estimated that about 2% of females meets the diagnostic criteria for antisocial personality disorder in comparison to 6% of males with the diagnosis (Trull et al., 2010). Little is also known about effects of psychopathic traits differences in factor expression on emotional processing characteristics in the general female population (Sutton, Vitale & Newman, 2002). It has been suggested however that primary and secondary variants of psychopathy in healthy females present distinct neural patterns of fear processing (Sethi et al., 2018) and manifest divergent psychophysiological reactivity to anger (Goulter et al., 2019). Furthermore, the emotional brain responses tend to be stronger in females than in males (Weisenbach et al. 2014).

### The Current Study

Brain activity patterns have been shown to predict various personality features such as negative-affect personality traits (Fernandes et al., 2017), narcissistic traits in healthy adults (Feng et al., 2018), as well as psychopathic traits in offenders (Sato et al., 2011) and in healthy individuals (Kimonis et al., 2006). Individual differences in personality traits, including psychopathic traits, also influence emotional processing in the brain (Hamann & Canli, 2004). Since movies are shown to provoke robust emotions in participants (Schaefer et al., 2010) and are widely used as naturalistic stimuli in neuroimaging studies (Jääskeläinen et al., 2021), we chose to examine how the perception of basic emotions (fear, sadness, happiness, anger, disgust, surprise; Ekman, 1992) in emotional movies is associated with the regional brain activity in female subjects with PP and SP traits. In this study, healthy female Finnish participants (n=50) filled Levenson Self-Report Psychopathy (LSRP) Scale that measures the level of primary and secondary psychopathy traits (Levenson et al., 1995). Then the subjects watched series of movie clips (n=39, duration 114 min.) during functional magnetic resonance imaging (fMRI) of brain activity. The movies were previously rated with separate group of participants (n=16) for the intensity of six basic emotions and neutral stimuli. In order to examine how individuals are similar in the neural responses to movies, we performed the pairwise intersubject correlation (ISC) of stimulus-induced brain activity. We then applied a whole brain univariate two-tailed t test to evaluate whether the mean ISC differed between the groups with low and high scores in primary and secondary psychopathy. Here we show that 1) primary and secondary psychopathic personality features in healthy females are linked with differential brain activity patterns to movies and 2) the differences in activation of the brain might be explained by emotional stimulation.

## METHODS AND MATERIALS

### Participants

Originally, fifty healthy Finnish-speaking right-handed female university students fluent in English participated in the fMRI study (age range: 20-40, mean age 24.9±4.4). Three participants were excluded from analyses due to excessive head motion (framewise displacement > 1.5 mm in >2 of 5 fMRI runs), hence data from 47 subjects were used in analysis. The study was approved by the Research Ethics Committee of Aalto University and all the subjects signed a written informed consent. The experiment lasted approximately 2 hours on two days, and participants received 20e/h as compensation.

### Psychopathic traits measure

Finnish-translated Levenson’s Self-Report Psychopathy (LSRP) Scale was used to evaluate the psychopathic personality traits. LSRP is a 26-item questionnaire with 5-point Likert scale (1-strongly disagree; 5-strongly agree) designed to distinguish between primary (16 questions) and secondary (10 questions) psychopathic features in noninstitutionalized populations (Levenson et al., 1995). The primary scale measures the manipulation, callousness, deceit and selfishness (affective-interpersonal dimension), whereas secondary scale measures antisocial behaviors with impulsivity and emotional disturbance (antisocial-behavioral dimension). The average scores in male population was shown to be 32.96 on PP scale and 20.04 on SP scale, while in female population 27.67 on PP scale and 19.03 on SP scale (Levenson et al., 1995).

In our participants group, the mean score was in PP scale 30.9±7 (max. score=53, min. score=20) and in SP scale 23±4 (max. score=30, min. score=14; see **Supplementary Figure S5)**. The PP scores were statistically different from normal distribution (Anderson-Darling normality test, p=0.04) and the SP scores were normally distributed (p=0.3) (see **Supplementary Figure S4).** The scores on PP and SP scales were weakly correlated (Spearman correlation coefficient, r=0.05, p=0.7). The participants were grouped into low PP/SP and high PP/SP groups based on the upper (>75%) and lower (<25%) quartiles on the PP and SP scale.

### fMRI Stimulus

Originally, a total of 54 movie clips from Hollywood movies previously shown to elicit strong emotions were selected from the database created by Schaefer et al. (2010). We excluded black-and-white films to minimize low-level visual differences between movie scenes. Eventually, for the final set we chose 39 movie clips (length 0:16-6:57; total duration 114 minutes, see **Supplementary Table S2**). Prior to data collection, the 13 Finnish-speaking female participants from the same population as fMRI subjects, rated the intensity of 63 different emotions (in English; Likert scale from 0 to 4) in the split movie scenes in order to divide the movie stimuli into the running sessions (*runs*) with roughly similar emotional content (for a list of emotions and selection process see **Supplementary Figure S1** and **Table S1**). Based on the ratings, we calculated the average intensity of each of 63 emotion categories for each movie scene by averaging across the scene-wise clips. We then arranged movies by the most intense emotion categories they elicited, and manually selected the scenes to cover a wide and equal range of different emotion categories. This selection procedure led to dividing an fMRI movie stimuli into five *runs* of similar emotional content as defined by the pilot ratings (8 scenes in runs 1-4 and 7 scenes in run 5; for detailed information see **Supplementary Methods and Materials**).

### Ratings of emotions in the movies

To generate final emotional content for the selected 39 movie clips, another 16 independent Finnish-speaking female raters from the same population as fMRI subjects, evaluated the perceived and experienced emotional content of the clips, cut into 3-10 s long scenes based on original cut points of the movie and quiet periods in the soundtrack. Using an in-house developed web tool (in Finnish; **Supplementary Figure S3**) the raters had to choose the level of intensity of each of 63 emotions elicited by the scene using a Likert scale from 0 to 4. The participants rated the movies in a random order, while the order of the scenes was retained. The participants were encouraged to rate all scenes of a movie one after another to enable the rating of slowly developing emotions (if participants were idle for a long period, a popup appeared to suggest that they restart the movie from the first scene). However, the participants were also encouraged to have breaks between movies to avoid fatigue during the long rating process. The raters were also compensated for their effort.

### Experimental design

The fMRI participants filled LSRPS online before the first fMRI session. During fMRI, subjects watched the 39 short movies divided into five runs (duration 21-24 min per run, 8-9 clips per run, total duration 114 minutes) and split into two fMRI sessions conducted on separate days. The order of runs was counterbalanced across participants, however the order of trials within a run was the same for all. A trial started with a fixation cross presented for 9.6 seconds, followed by a movie clip. After the last clip, a fixation cross appeared again for 9.6 seconds. The participants were instructed to watch the movies attentively. During the experiment, participants’ alertness was monitored with eye-tracking and face camera. In addition to fMRI, 10-min resting state and 6-min anatomical scans were obtained. After both fMRI sessions subjects were debriefed and asked which movies they recognized.

### Stimulus presentation and physiological measures

The sound track of the movies was delivered through Sensimetrics S14 insert earphones (Sensimetrics Corporation, Malden, Massachusetts, USA). The visual stimuli were back-projected on a semitransparent screen using a Panasonic PT-DZ110XEJ data projector (Panasonic, Osaka, Japan) and from there viewed by the participant via a mirror fixed to the head coil. Stimulus presentation was controlled with Presentation software (Neurobehavioral Systems Inc., Albany, CA, USA). Heart rate (finger pulse transducer) and respiration (respiratory belt) data was collected with BIOPAC MP150 Data Acquisition System (BIOPAC System, Inc., USA). For details see **Supplementary Methods and Materials.**

### MRI data acquisition and processing

MRI data were collected using Siemens Skyra 3T MRI with 30-channel head coil at the Advanced Magnetic Imaging Centre, Aalto Neuroimaging, Aalto University. High-resolution anatomical images (1 x 1 x 1 mm voxel size) were acquired using T1-weighted MP-RAGE sequence with 2530-ms TR. Functional MRI was obtained with whole-head T2*-weighted echo-planar imaging pulse sequence (TR=2400 ms, TE=24 ms, flip angle=70°, and 3 x 3 x 3 mm resolution; fat-suppression with bipolar water excitation radio frequency pulse). fMRI data were preprocessed using FMRIPREP (v1.3.0) (Gorgolewski et al., 2011; Esteban et al., 2019). Each T1-weighted volume was corrected for intensity non-uniformity using N4BiasFieldCorrection (v2.1.0) and skull-stripped using ANTs (v2.1.0; the OASIS template). Brain surfaces were reconstructed using *recon-all* from FreeSurfer (v6.0.1) which also produced volume estimates of cortical (34 for each hemisphere) and subcortical (63) brain structures for each subject. For details see **Supplementary Methods and Materials.**

### Data preparation and group comparisons

First, we checked that psychopathy scores had sufficient variance across the subjects (see **Supplementary Figure S4**) and the emotional ratings had high inter-rater agreement (see **Supplementary Figure S10**). From 63 emotions we selected 6 perceived basic emotion and neutral stimuli that had been most relevant in psychopathy personality research. The fMRI data consisted of the concatenated blood oxygenation level dependent (BOLD) activity over the total length of all movies taken together to evaluate the effects over the entire experiment. To evaluate how the effects related to specific basic emotions, we repeated the analyses with concatenated BOLD data over only those timepoints that received non-zero emotional ratings for each perceived emotion. First, we calculated the pairwise intersubject correlation (ISC) matrices between all pairs of participants. To compare the difference in ISC between participants scoring high vs. low on psychopathic traits, we extracted pairwise ISC values from participants in the upper (>75%) and lower (<25%) quartiles on the PP and SP scale. We then applied a whole brain univariate two-tailed t test to evaluate at each voxel, whether the mean ISC differed between the groups. Statistical significance was evaluated with element-wise permutation of the correlation matrix. The outcome brain maps were cluster corrected with p value threshold 0.0001. To summarize the data, we searched for the local maxima in the significant clusters (min. size 105 voxels) showing group differences. These cluster maxima were named based on the atlas labels of the Brainnetome atlas (Fan et al.,2016), the Probabilistic atlas of the human cerebellum (Diedrichsen et al., 2009; Diedrichsen et al., 2011) and the Brainstem Navigator tool (Bianciardi, 2021). Additionally, we calculated the percentage of volume of each of the atlas regions that showed significant group differences and summarized this data for the major lobes of the brain with total 334 structures.

## RESULTS

### Psychopathic traits and neural responses to movies

**Figure 1** presents the group differences of intersubject correlation (ISC) between participant scoring low vs. high in primary (PP) and secondary (SP) psychopathy. The box plots indicate the percent of the area within brain regions that showed significant group differences in ISC of neural responses over all the significant clusters. The locations of the significant clusters are shown on the brain maps.

**Figure 1.**
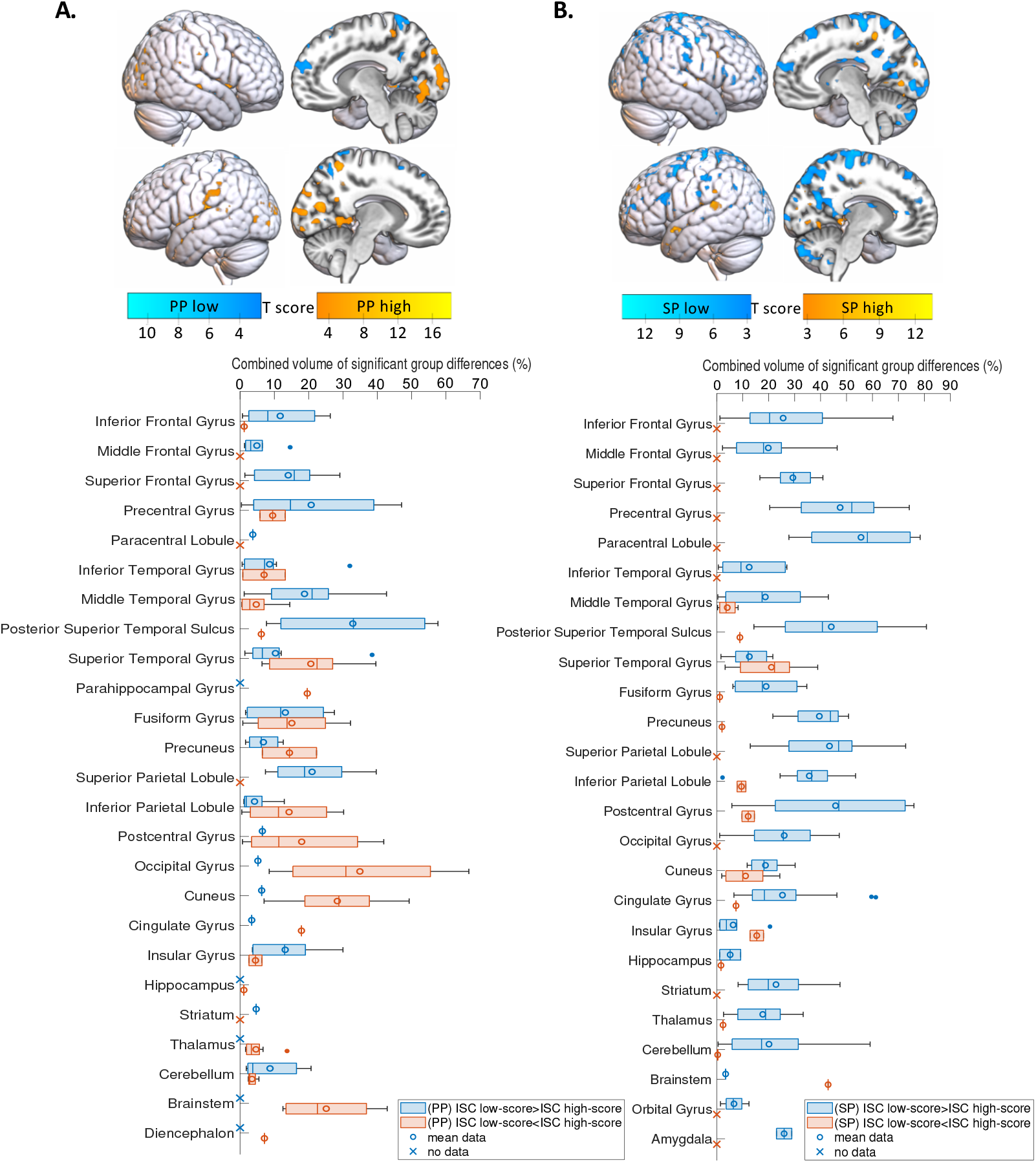
Percentage of volume within the brain regions with significant group differences between subjects with primary (PP) and secondary (SP) psychopathy scores during watching the movie stimuli. The box plots (down) show median and average percentage of the regions’ volumes based on intersubject correlation (ISC) similarity between the participants and their score in PP (A) and SP (B). The boxes indicate the interquartile range and the whiskers the maximum and minimum values over significant clusters. Regions with no significant effects have been excluded. The brain maps (up) show the brain activation areas similar in participants with low and high psychopathy score.

Individuals scoring low vs. high on PP showed higher ISC in the Posterior Superior Temporal Sulcus (pSTS; approximately 35% of volume), Precentral Gyrus (preCG; 20%), Superior Parietal Lobule (SPL; 20%), Middle Temporal Gyrus (MTG; 20%), Insular Gyrus (15%), cerebellum (10%) and striatum (5%). Individuals with high vs. low PP score exhibited higher ISC in the Occipital Gyrus (OG) with approximately 35% of significant volume within the region, as well as in Cuneus (30%) and brainstem (30%).

The participants with low score in SP showed higher ISC than those with high SP scores in the Paracentral Lobule (PCL), with approximately 60% of the region showing a significant group difference, preCG (50%), Postcentral Gyrus (45%), pSTS (45%), striatum and cerebellum (20%). Additional effects were observed in parts of the amygdala (25%) and the striatum (25%). The ISC of the subjects with high vs. low SP sores was significantly higher in only a few regions including parts of the brainstem (45%), Superior Temporal Gyrus (STG; 20%) and Insular Gyrus (15%).

**Figure 2** show the proportional volume within main areas of the brain that was significantly different between ISC neural responses of participants with low and high PP/SP score while watching the films. The highest proportion of effects in subjects with low PP score was found mostly in temporal areas (19% activation comparing to other regions) whereas in participant with high PP score the proportional effects were high mostly in occipital areas (23%) and brainstem (18%). The participants with low score in SP had proportionally highest effects in frontoparietal areas (21%) and subjects with high SP score showed high brainstem effects within the region (43%) in comparison to other structures.

**Figure 2.**
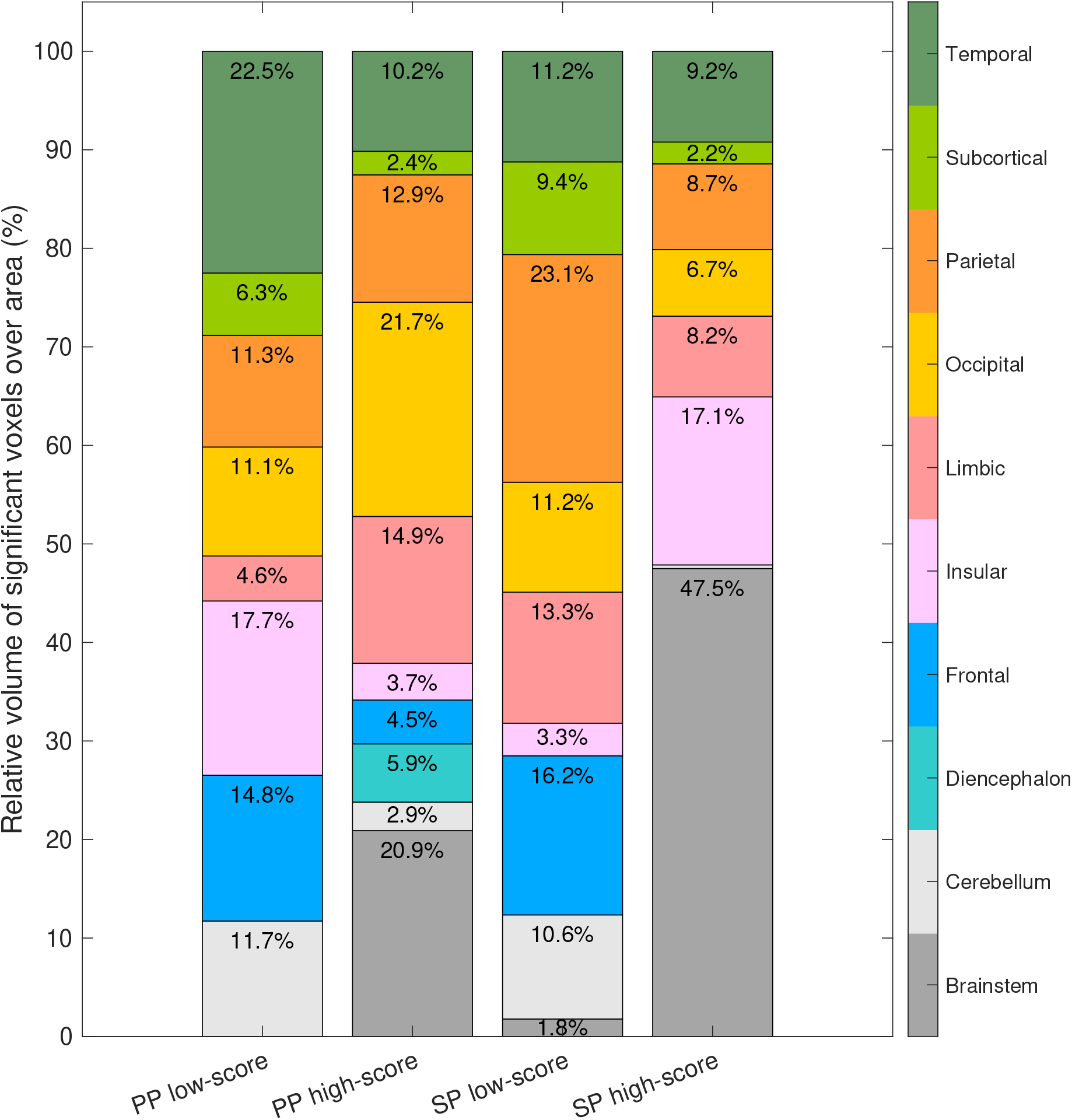
Mean percentage of volume within the main brain areas with significant ISC group differences between subjects with primary (PP) and secondary (SP) psychopathy scores. The brain regions were divided to show the significant group differences within the main brain areas in low/high PP and SP score. The relative volume of significant voxels in each brain area is shown in proportion to the total significant volume over the entire brain (summarized to 100%).

### Participants’ psychopathy score and neural reaction to basic emotions in the movies

In addition to neural response in participants with psychopathy traits to films as a whole, we investigated whether the perceived emotional stimuli involving six basic emotions present in the movies may explain the groups differences in regional activity. We found that in subjects with low PP score the perception of fear predicted closely the increases of ISC over pSTS and MTG seen towards all movies. Differences in ISC in the cerebellum and insula were predicted by negative emotions, mostly by perception of fear, anger and disgust, and striatum was activated mostly by the perceived anger (**Figure 3A)**. In individuals with high PP score the differences in ISC in OG and brainstem were predicted by perception of fear and surprise and in Cuneus by perception of fear and anger. The ISC differences in insula was predicted by perceiving the neutral stimuli (**Figure 3B).**

**Figure 3.**
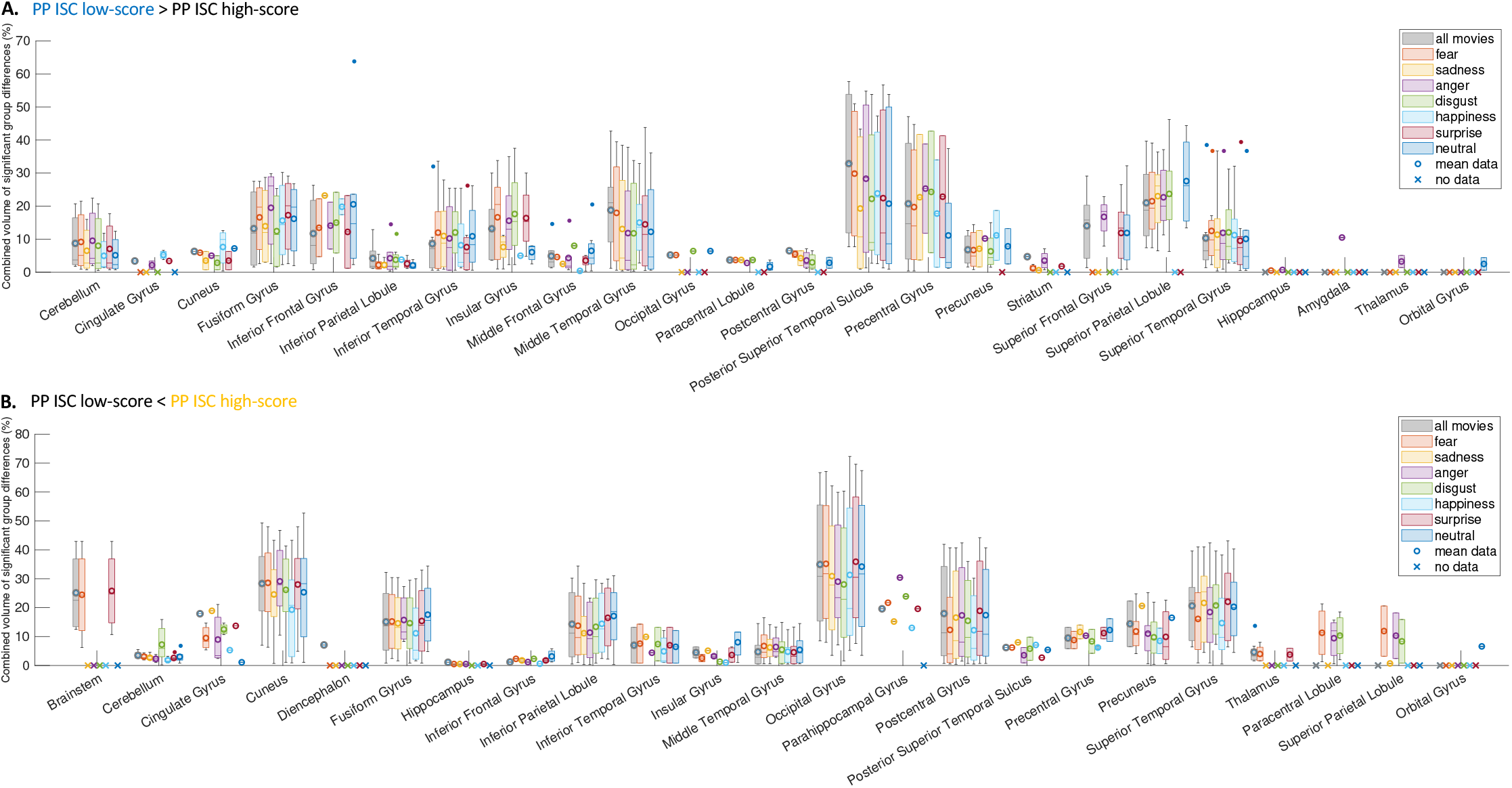
Percentage of volume within the brain regions with significant group differences between subjects with primary psychopathy (PP) according to perception of emotions. The box plots show median and average percentage of the regions’ volumes based on intersubject correlation (ISC) similarity between the participants with low PP score (A) and high PP score (B) in response to perception of 6 basic emotions. The boxes indicate the interquartile range and the whiskers the maximum and minimum values over significant clusters. Regions with no significant effects have been excluded. The grey bar as the significant group difference to all movies is added for comparison.

In general, in individuals with lower vs. higher SP scores, the brain regions that showed highest ISC increases in response to all movies (PCL, preCG, postCG, pSTS) were best predicted by perceived fear and anger (**Figure 4A**). The effects in amygdala, cerebellum, Cingulate Gyrus (CG), insula and striatum were well predicted by the perception of negatively valenced affect, predominantly fear, anger and surprise. On the other hand, the brainstem effects in participants with high SP score was predicted by the perception of fear and insula by perception of sadness. The STG showed highest differences during perceived fear and sadness. The Cuneus and CG differences were similar to the mean effects during all emotions (**Figure 4B**).

**Figure 4.**
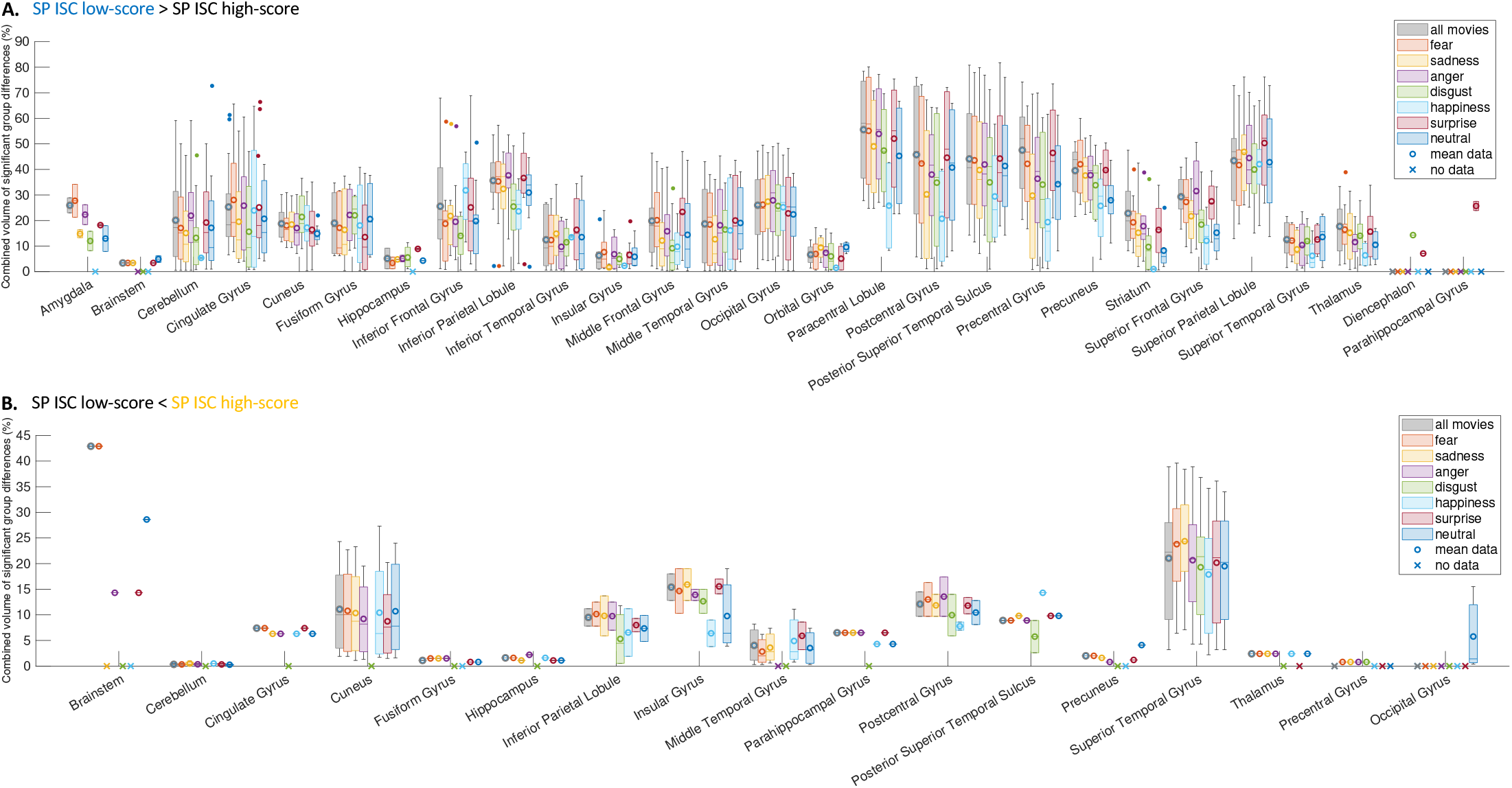
Percentage of volume within the brain regions with significant group differences between subjects with secondary psychopathy (SP) according to perception of emotions. The box plots show median and average percentage of the regions’ volumes based on intersubject correlation (ISC) similarity between the participants with low SP score (A) and high SP score (B) in response to perception of 6 basic emotions. The boxes indicate the interquartile range and the whiskers the maximum and minimum values over significant clusters. Regions with no significant effects have been excluded. The grey bar as the significant group difference to all movies is added for comparison.

## DISCUSSION

Our main finding was that neural responses to emotional movies depend on primary (PP) and secondary psychopathy (SP) traits in healthy women sampled from the general population. Our results generally align with previous findings showing that separate psychopathy personality types are linked with differential brain activity patterns. We suggest that already normal variability in psychopathic traits in the typical population is manifested in differential neural responses to movies. The emotion-specific analyses suggest that these differences are also modulated by emotional processing.

Neural activity in participants with PP traits during watching movies showed that subjects with low score were more similar in fronto-temporal areas and cerebellum whereas subjects with high score showed higher similarity in occipital regions and brainstem (compare **Figure 1A** and **Figure 2**). These cortical regions are linked with audiovisual processing that include monitoring of social information and identification of emotional stimuli (Dermody et al., 2016). The simultaneous engagement of sensorimotor and subcortical systems during emotional stimulation such as watching a movie is associated with vicarious representation of others’ emotional states (Decety et al., 2013; Marcoux et al., 2013). The striatal and insular activity to movies in low PP was related to more consistent striatal and insular responses during anger and fear perception (see **Figure 3A**), suggesting that subjects with low PP traits may show heightened sensitivity to thread and aggression-related affect. Even though the activity in striatum and insula has been generally connected to pleasure experience and reward-directed action (Pizzagalli et al., 2009), it was also indicated in promoting aversive reaction and avoidance behavior as response to negative stimulation (Volman et al., 2013; Zeki & Romaya, 2008).

We found high PP traits to be associated with low insular response and reduction of striatal engagement. The psychopathy was connected previously with abnormal striatal and insular activity during viewing movies (Nummenmaa et al., 2021) and increased task-related activity in striatum (Deming & Koenigs, 2020) which contradicts our results. Nevertheless, the Decety et al. (2013) found that the absence of coherent striatal activation was a sign of lack of vicarious response while engaging sufficient sensorimotor resonance during mentalizing about others’ emotions, which could be consistent with our findings. The disengagement of striatum while watching movies could also lead towards diminished sensitivity to emotional stimuli, particularly to negative and unpleasant emotional states as previously found in psychopathy traits (Verona, Patrick & Joiner, 2001; Kimonis et al., 2008). However, the simultaneously heightened similarity of activity of Cingulate Gyrus (CG) and brainstem that we found related to the perception of sadness and fear respectively (see **Figure 3B**) challenge this assumption. The CG mediates emotion and memory, and process especially threat-related emotional information (Rolls, 2019). The brainstem has also been increasingly reported to be involved in emotional and empathetic arousal (Price & Drevets, 2010). Therefore, despite the reports of diminished consistency of activity of brainstem and autonomic nervous system in psychopathy (Marsh, 2013) it is possible that enhanced autonomic arousal towards negative stimuli is a feature of healthy participants with high traits in PP.

In subjects with low SP traits, we found wide-spread increases in the similarity of activations across the brain, mostly in fronto-parietal areas and cerebellum (**Figure 2)**. Similarly to low PP, this increased similarity may indicate emotional mentalizing and sensorimotor resonance with movie characters. However, low SP group also showed heightened similarity in activation of striatum and CG as well as in Insular Gyrus, Thalamus and Amygdala (see **Figure 4A**). These regional increases, and particularly amygdalar response, were also dominant during the perception of fear. Since all those structures are strongly implicated in production and regulation of fear and anxiety (Huggins et al., 2021), the response of fear-related neurocircuitry suggests enhanced sensitivity to anxious stimuli in low SP comparing to low PP. The SP type have been generally reported to originate from early life trauma and be related to abuse-connected responses and emotionality (Skeem et al., 2007). Thus, potentially the presence of SP traits, even on low-end continuum, might manifest in heightened responsivity to negative and threat-related emotions.

Furthermore, failure to recognize others’ fear or to generate empathic activation in the amygdala is a hallmark of primary psychopathy (Blair, 2006) and as expected, we did not find amygdalar response in individuals with high PP traits. We also have not found the prominent subcortical similarity in response to threat in people with high SP traits which could confirm that the experience of fear and anxiety in SP might be intact (Hofmann, Schneider & Mokros, 2021) or diminished (Shultz et al., 2016). On the other hand, we found the high scoring SP subjects similar in insular response to perceiving negative emotions what contradicts the previous findings as psychopathy was generally associated with decrease of insula activity during watching fear of others (Deming et al., 2020). Furthermore, we discovered the profoundly greater similarity in activation of brainstem to the perception of fear in high SP when comparing to high PP traits (see **Figure 4B**). The brainstem has been shown previously to influence the generation and experience of emotions by modulating the arousal, autonomic function, motor control and somatosensation (Venkatraman et al., 2017). This suggests that high SP traits might be linked with greater subcortical brainstem-dependent arousal to fearful stimuli. It also further indicates that SP traits manifest in higher general sensitivity to negative emotions and anxiety when comparing to high scoring PP individuals (Lee & Salekin, 2010; Sethi et al., 2018).

## LIMITATIONS

First, our subject group included only young females. Thus the generalization of the results to other populations might be restricted. Second, the agreement between the raters of the emotional content of the movies varied across emotions (see **Supplementary Figure S10**), hence theoretically particular emotional results in the brain may be less reliable. Moreover, there were two separate groups of subjects that rated the emotions in the movies and watched the movies in fMRI. Future studies could replicate the method with basic emotions not only in perception but also experience of affect, and include higher number of raters in order to elevate the inter-raters reliability (IRR). Third, the low variability in secondary psychopathy scores among the participants could partly influence the limited associations with hemodynamics and SP scores (see **Supplementary Figure S4** and **S5**). However, percentile split in t test that we used should decrease this effect. Thus it is probable that neural activity in subjects with high SP score showed greater idiosyncrasy. In the future, this problem could be omitted by stricter preselection of the subjects manifesting low and high personality traits of both types of psychopathy.

## CONCLUSIONS

We conclude that neural responses to movies depend on primary and secondary psychopathy traits in the general population and the emotional stimulation can further improve the understanding of the neural differences in psychopathy traits. We found that participants low on primary and secondary psychopathy traits showed higher cortical and subcortical activity when watching emotional movies, which possibly indicates the somato-motoric simulation and reflecting the emotions of movie characters. Furthermore, the individuals low on SP traits showed heightened consistency of striatal and subcortical reactivity predominantly in observing fearful content in the movies. Thus, participants with any level of SP traits may possibly manifest higher sensitivity towards negatively-valenced affect. On the other hand, the subjects with high primary and secondary psychopathy traits presented a lack of striatal engagement while activating the cortical regions, which is consistent with reduced vicarious engagement to others’ emotional states. Moreover, the participants scoring high on SP scale showed increased response in insula and brainstem to negative emotions and particularly fear when comparing to high PP score. Thus, this suggest that high SP traits are more connected with potent autonomic and subcortical response while watching threat-related stimuli.

