## Supplemental Information for "Psychopathic traits predict neural responses to emotional movies in the general population"

### 1. Supplementary Methods and Materials

#### 1.1. FMRI stimulus

**a) Selection of emotion categories**

The emotion categories were selected based on previous literature and on similarity ratings collected from a separate set of participants. First, literature introducing multiple emotion categories (Adolphs, 2002; Cowen & Keltner, 2017; Saarimäki et al., 2018; Skerry & Saxe, 2016) was searched for an initial list of categories (Step 1). Second, to ensure that the selected emotion categories would cover the whole emotion space, we added the emotion categories from Shaver et al. (2001). This resulted in a list of 128 categories (Step 2). Third, as the participants of the study were Finnish, the emotion categories were translated to Finnish using a dictionary of Finnish emotion words (Tuovila, 2005). Fourth, we collected similarity ratings of these 128 emotion categories to choose the categories that would complement the list chosen in Step 1. Similarity ratings were collected from a sample of twenty female participants that did not participant in the other behavioral or fMRI experiments of the current study. The similarity ratings were collected using an online tool (https://version.aalto.fi/gitlab/eglerean/sensations) where a list of all emotion categories is presented next to a large canvas. Using a mouse, the participants arranged the emotion categories on the canvas so that categories that feel more similar were placed closer to each other. The results of the average similarity ratings for the 128 emotions displaying the pairwise distance between emotion categories are shown in **Figure S1**. From the similarity ratings, we extracted 8 clusters of emotion categories. Further, we divided the category *pain* to two categories, one for mental pain and one for physical pain. Thus, the final list of 63 emotion categories consisted of 1) the categories listed in Step 1, and 2) the categories that complemented the emotion space based on visual inspection (Step 4, see **Table S1**), thus included: *admiration, adoration, amusement, anger, annoyance, anxiety, apprehension, awe, awkwardness, boredom, calmness, confusion, contempt, contentment, craving, defeat, despair, devastation, disappointment, disgust, dislike, displeasure, embarrassment, emotional arousal, emotional pain, entrancement, excitement, fear, fury, glumness, gratitude, grief, guilt, happiness, hope, horror, hostility, humiliation, hurt, impressed, infatuation, interest, insecurity, jealousy, joy, loneliness, longing for, love, neutral, nostalgia, physical pain, pity, pride, relief, romance, sadness, satisfaction, sexual desire, shame, suffering, surprise, thrill,* and *zeal.*

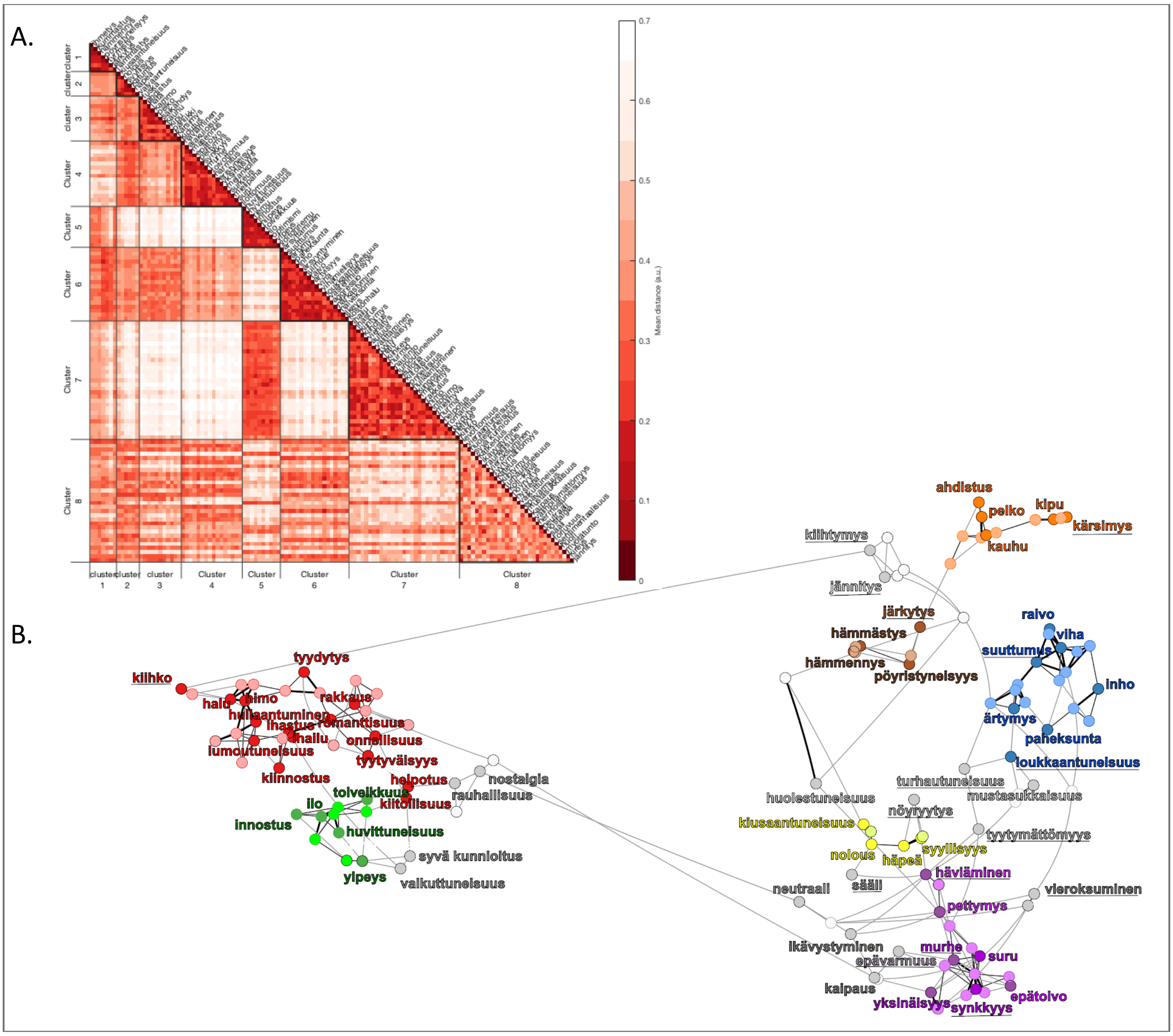


**Figure S1.** Similarity ratings for the 128 emotion categories. A. Clustering of emotion categories. B. Network of emotion categories with colors denoting the clusters. Note that Cluster 8 (in grey) is sparse and includes categories not belonging to other clusters. Labels are shown for the categories included in the final list of 63 emotions. The emotion categories added based on the similarity ratings are underlined. The categories excluded from the final list are shown with white dots without a label.

**Table S1.** Final list of emotion categories. References: Cowen & Keltner 2017; Skerry & Saxe 2016; Saarimäki et al. 2018; Adolphs 2002; Shaver et al. 2001

| Cluster | Emotion category (FIN) | Emotion category (ENG) | References | Inclusion step |
|---|---|---|---|---|
| **1** | hämmennys | confusion | 1 | 1 |
| **1** | järkytys | devastation | 2 | 1 |
| **1** | hämmästys | surprise | 1-5 | 1 |
| **2** | kiusaantuneisuus | awkwardness | 1 | 1 |
| **2** | nolous | embarrassment | 2, 4, 5 | 1 |
| **2** | syyllisyys | guilt | 2-5 | 1 |
| **2** | häpeä | shame | 2, 5 | 1 |
| **3** | ahdistus | anxiety | 1, 5 | 1 |
| **3** | pelko | fear | 1, 3-5 | 1 |
| **3** | kauhu | horror | 1, 5 | 1 |
| **3** | kipu (fyysinen) | pain (physical) | 1 | 1* |
| **3** | kipu (henkinen) | pain (mental) | 1 | 1* |
| **3** | kärsimys | suffering | 5 | 4 |
| **4** | lannistuneisuus | defeat | 5 | 4 |
| **4** | epätoivo | despair | 3, 5 | 1 |
| **4** | pettymys | disappointment | 2, 5 | 1 |
| **4** | synkkyys | glumness | 5 | 4 |
| **4** | murhe | grief | 5 | 4 |
| **4** | yksinäisyys | loneliness | 2, 5 | 1 |
| **4** | suru | sadness | 1, 3-5 | 1 |
| **5** | huvittuneisuus | amusement | 1, 5 | 1 |
| **5** | innostus | excitement | 1, 2, 5 | 1 |
| **5** | toiveikkuus | hope | 2, 5 | 1 |
| **5** | ilo | joy | 1, 2, 5 | 1 |
| **5** | ylpeys | pride | 2-5 | 1 |
| **6** | suuttumus | anger | 1-5 | 1 |
| **6** | ärtymys | annoyance | 2, 5 | 1 |
| **6** | halveksunta | contempt | 3-5 | 1 |
| **6** | inho | disgust | 1-5 | 1 |
| **6** | raivo | fury | 2, 5 | 1 |
| **6** | viha | hostility | 5 | 4 |
| **6** | loukkaantuneisuus | hurt | 5 | 4 |
| **7** | ihailu | admiration | 1, 4 | 1 |
| **7** | ihastus | adoration | 1, 4 | 1 |
| **7** | tyytyväisyys | contentment | 2, 5 | 1 |
| **7** | seksuaalinen halu | sexual desire | 1 | 1 |
| **7** | lumoutuneisuus | entrancement | 1 | 1 |
| **7** | kiitollisuus | gratitude | 2, 3 | 1 |
| **7** | onnellisuus | happiness | 3-5 | 1 |
| **7** | hullaantuminen | infatuation | 4, 5 | 1 |
| **7** | kiinnostus | interest | 1 | 1 |
| **7** | rakkaus | love | 2, 4, 5 | 1 |
| **7** | himo (ei seksuaalinen) | craving | 1 | 1 |
| **7** | helpotus | relief | 1, 5 | 1 |
| **7** | romanttisuus | romance | 1 | 1 |
| **7** | tyydyttyneisyys | satisfaction | 1 | 1 |
| **7** | kiihko | zeal | 5 | 4 |
| **8** | huolestuneisuus | apprehension | 2, 5 | 1 |
| **8** | syvä kunnoitus | awe | 1 | 1 |
| **8** | ikävystyminen | boredom | 1 | 1 |
| **8** | rauhallisuus | calmness | 1 | 1 |
| **8** | vieroksuminen | dislike | 5 | 4 |
| **8** | tyytymättömyys | displeasure | 5 | 4 |
| **8** | kiihtymys | excitement | 5 | 4 |
| **8** | nöyryytys | humiliation | 5 | 4 |
| **8** | vaikuttuneisuus | impression | 2 | 1 |
| **8** | epävarmuus | insecurity | 5 | 4 |
| **8** | mustasukkaisuus | jealousy | 2, 5 | 1 |
| **8** | kaipaus | longing for | 3, 5 | 1 |
| **8** | neutraali | neutral | 3 | 1 |
| **8** | nostalgia | nostalgia | 1, 2 | 1 |
| **8** | sääli | pity | 5 | 4 |
| **8** | jännittävyys | thrill | 5 | 4 |

**b) Selection of movie clips**

The sets of movie clips (54) were selected from existing emotion-elicitation database that included in majority Hollywood movies (Schaefer et al., 2010). We excluded black-and-white films to minimize low-level visual differences between movie scenes and we included only one movie clip from the same movie. The movie clips were then cut into 3-10 seconds long scenes based on the original cut-points preferring local loudness minima in the soundtrack of the movies to avoid splitting the videos in the middle of continuous actions. Where the time between subsequent cuts exceeded the maximum clip duration of 10 seconds, the cut interval was first equally split into the minimum number of <10-second windows. The splitting points were then adjusted to coincide with the closest loudness minima of the soundtrack. The 13 Finnish-speaking female participants from the same population as fMRI subjects, rated the intensity of 63 different emotions (in English; Likert scale from 0 to 4) in the split movie scenes in order to divide the movie stimuli into the *runs* with roughly similar emotional content. Therefore from the ratings, we calculated the average intensity of each of 63 emotion categories for each movie scene by averaging across the scene-wise clips. We selected the emotions with highest averages across movies and shared them into negative, positive and neutral/confusing emotional content (**Figure S2**). We then chose 9 emotion categories according to valence and highest averaged ratings, and arranged movies by the most intense emotion categories they elicited. We manually selected the scenes to cover a wide and equal range of different emotion categories. This selection procedure led to a final set of 39 movie clips (length 0:16-6:57; total duration 114 minutes, **Table S2**) which were divided in the fMRI study into five running sessions (*runs*) of similar emotional content as defined by the pilot ratings (8 scenes in runs 1-4 and 7 scenes in run 5; **Figure S3**).

### Negative emotions

**Fear & Anxiety**

| 16 | 18 | 28 | 38 | 46 | 55 |
|---|---|---|---|---|---|
| 'anxiety' | 'fear' | 'anxiety' | 'anxiety' | 'anxiety' | 'anxiety' |
| 3.2 | 1.8 | 3.0 | 1.3 | 2.2 | 2.3 |
| 6 | 5 | 6 | 5 | 6 | 6 |
| 'fear' | 'anxiety' | 'fear' | 'fear' | 'fear' | 'fear' |
| 2.7 | 1.7 | 2.6 | 1.2 | 1.8 | 1.9 |
| 6 | 5 | 6 | 6 | 5 | 6 |
| 'concerned' | 'concerned' | 'concerned' | 'concerned' | 'confusion' | 'unsure' |
| 1.4 | 1.4 | 1.1 | 0.9 | 0.8 | 0.7 |
| 4 | 4 | 3 | 3 | 3 | 3 |
| 'despair' | 'sadness' | 'despair' | 'surprise' | 'concerned' | 'concerned' |
| 0.9 | 1.3 | 0.9 | 0.7 | 0.7 | 0.6 |
| 3 | 6 | 4 | 4 | 3 | 3 |
| 'hostile' | 'interested' | 'agitated' | 'neutral' | 'anger' | 'unfriendly' |
| 0.8 | 1.0 | 0.9 | 0.4 | 0.5 | 0.4 |
| 2 | 2 | 3 | 4 | 2 | 1 |

**Disgust**

| 7 | 22 | 31 | 35 | 57 |
|---|---|---|---|---|
| 'disgust' | 'disgust' | 'disgust' | 'disgust' | 'disgust' |
| 2.8 | 1.4 | 1.5 | 2.6 | 3.2 |
| 6 | 6 | 6 | | 6 |
| 'anxiety' | 'anxiety' | 'interested' | 'confusion' | 'fear' |
| 2.2 | 0.9 | 1.1 | 1.1 | 1.3 |
| 6 | 4 | 3 | | 5 |
| 'fear' | 'interested' | 'fear' | 'surprise' | 'confusion' |
| 1.2 | 0.6 | 0.7 | 0.4 | 1.0 |
| 4 | 3 | 3 | | 4 |
| 'confusion' | 'neutral' | 'anxiety' | 'puzzlement' | 'anxiety' |
| 0.4 | 0.4 | 0.7 | 0.4 | 0.8 |
| 2 | 5 | 4 | | 2 |
| 'hurt' | 'sadness' | 'concerned' | 'anxiety' | 'surprise' |
| 0.4 | 0.4 | 0.5 | 0.4 | 0.5 |
| 1 | 3 | 3 | | 2 |

**Pain**

| 10 | 21 | 50 | 27 |
|---|---|---|---|
| 'sadness' | 'sadness' | 'anxiety' | 'anxiety' |
| 2.2 | 2.1 | 1.8 | 2.0 |
| 6 | 6 | 6 | 6 |
| 'anxiety' | 'fear' | 'fear' | 'sadness' |
| 2.0 | 1.7 | 1.4 | 1.7 |
| 6 | 5 | 5 | 5 |
| 'fear' | 'pain' | 'concerned' | 'pain' |
| 1.5 | 1.4 | 1.2 | 1.3 |
| 6 | 4 | 3 | 4 |
| 'pain' | 'anxiety' | 'sadness' | 'concerned' |
| 1.0 | 1.4 | 0.5 | 1.2 |
| 3 | 5 | 2 | 4 |
| 'despair' | disagreement | 'pain' | 'fear' |
| 0.9 | 1.2 | 0.4 | 1.1 |
| 3 | 2 | 5 | 4 |

**Sadness**

| 9 | 36 | 43 | 52 | 62 |
|---|---|---|---|---|
| 'interested' | 'sadness' | 'sadness' | 'sadness' | 'neutral' |
| 1.7 | 1.8 | 1.3 | 1.5 | 1.3 |
| 5 | 6 | 6 | 6 | 6 |
| 'sadness' | 'anxiety' | 'anxiety' | 'neutral' | 'interested' |
| 1.4 | 1.1 | 1.2 | 1.0 | 1.2 |
| 5 | 6 | 6 | 6 | 5 |
| 'touched' | 'sorry' | 'fear' | 'sorry' | 'sadness' |
| 1.1 | 0.8 | 0.9 | 0.9 | 0.9 |
| 5 | 3 | 5 | 4 | 5 |
| 'love' | 'concerned' | 'unsure' | 'interested' | 'touched' |
| 0.6 | 0.7 | 0.8 | 0.6 | 0.6 |
| 1 | 4 | 4 | 3 | 6 |
| 'kind' | 'fear' | 'concerned' | 'anxiety' | 'anxiety' |
| 0.6 | 0.7 | 0.6 | 0.5 | 0.5 |
| 1 | 4 | 3 | 2 | 2 |

**Unfriendliness & possible Anger**

| 25 | 30 | 32 | 39 | 51 |
|---|---|---|---|---|
| 'anxiety' | 'anxiety' | 'anxiety' | 'fear' | 'anger' |
| 1.6 | 1.6 | 2.3 | 2.7 | 0.9 |
| 6 | 5 | 6 | 6 | 4 |
| 'fear' | 'unfriendly' | 'fear' | 'anxiety' | 'annoyed' |
| 1.2 | 1.0 | 1.7 | 1.9 | 0.8 |
| 5 | 3 | 5 | 5 | 3 |
| 'disgust' | 'fear' | 'concerned' | 'concerned' | 'irritated' |
| 0.8 | 1.0 | 1.1 | 1.8 | 0.8 |
| 4 | 4 | 3 | 4 | 4 |
| 'anger' | 'sadness' | 'interested' | 'anger' | 'anxiety' |
| 0.6 | 0.8 | 0.8 | 1.5 | 0.8 |
| 5 | 4 | 3 | 4 | 6 |
| 'confusion' | 'confusion' | 'anger' | 'sadness' | 'confusion' |
| 0.6 | 0.7 | 0.5 | 0.9 | 0.6 |
| 4 | 4 | 1 | 4 | 4 |

### Positive emotions

**Funny stuff**

| 14 | 23 | 56 | 61 |
|---|---|---|---|
| 'amusement' | 'interested' | 'amusement' | 'amusement' |
| 1.0 | 1.5 | 1.9 | 1.1 |
| 5 | 4 | 6 | 5 |
| 'interested' | 'laughter' | 'interested' | 'laughter' |
| 0.9 | 1.3 | 1.1 | 1.0 |
| 4 | 4 | 6 | 4 |
| 'neutral' | 'amusement' | 'happiness' | 'surprise' |
| 0.9 | 1.0 | 0.9 | 0.8 |
| 6 | 4 | 6 | 5 |
| 'laughter' | 'neutral' | 'laughter' | 'neutral' |
| 0.6 | 0.9 | 0.8 | 0.6 |
| 4 | 4 | 3 | 6 |
| 'annoyed' | 'confusion' | 'confusion' | 'confusion' |
| 0.4 | 0.9 | 0.8 | 0.6 |
| 2 | 3 | 5 | 4 |

**Moving stuff**

| 15 | 26 | 33 | 48 | 59 |
|---|---|---|---|---|
| 'touched' | 'neutral' | 'tenderness' | 'touched' | 'touched' |
| 1.2 | 1.4 | 1.8 | 1.2 | 1.3 |
| 6 | 3 | 4 | 4 | 5 |
| 'interested' | 'love' | 'romantic' | 'laughter' | 'interested' |
| 1.1 | 1.1 | 1.8 | 0.9 | 0.8 |
| 4 | 4 | 6 | 4 | 3 |
| 'happiness' | 'relaxed' | 'love' | 'amusement' | 'happiness' |
| 0.8 | 1.0 | 1.6 | 0.8 | 0.8 |
| 5 | 4 | 4 | 4 | 4 |
| 'love' | 'romantic' | easure (sensu | 'kind' | 'sadness' |
| 0.6 | 1.0 | 1.4 | 0.6 | 0.6 |
| 4 | 3 | 5 | 3 | 6 |
| 'tenderness' | 'tenderness' | 'touched' | 'love' | 'sorry' |
| 0.6 | 0.9 | 1.2 | 0.5 | 0.5 |
| 3 | 3 | 3 | 1 | 3 |

### Neutral and confusing

**Neutral stuff**

| 45 | 49 | 58 | 64 |
|---|---|---|---|
| 'neutral' | 'neutral' | 'neutral' | 'neutral' |
| 2.5 | 2.8 | 3.1 | 2.7 |
| 6 | 6 | 6 | 6 |
| 'relief' | 'interested' | 'interested' | 'confusion' |
| 0.4 | 0.3 | 0.6 | 0.3 |
| 1 | 1 | 4 | 2 |
| 'interested' | 'happiness' | 'unsure' | 'puzzlement' |
| 0.3 | 0.3 | 0.4 | 0.2 |
| 1 | 1 | 1 | 3 |
| 'bored' | 'pain' | 'puzzlement' | 'interested' |
| 0.2 | 0.0 | 0.2 | 0.2 |
| 1 | 0 | 2 | 1 |
| 'confusion' | 'disgust' | 'bored' | 'irritated' |
| 0.1 | 0.0 | 0.0 | 0.1 |
| 1 | 0 | 1 | 1 |

**Confusing stuff**

| 3 | 17 | 53 |
|---|---|---|
| 'anxiety' | 'unsure' | 'confusion' |
| 1.3 | 0.9 | 1.3 |
| 6 | 3 | 5 |
| 'confusion' | 'neutral' | 'disgust' |
| 1.1 | 0.8 | 0.8 |
| 5 | 4 | 4 |
| 'concerned' | 'confusion' | 'annoyed' |
| 1.0 | 0.8 | 0.8 |
| 4 | 4 | 3 |
| 'interested' | 'puzzlement' | 'bothered' |
| 1.0 | 0.5 | 0.6 |
| 4 | 5 | 2 |
| 'unsure' | 'disgust' | 'laughter' |
| 0.6 | 0.5 | 0.4 |
| 3 | 6 | 2 |

**Figure S2.** Division into valence of most intensive (out of 63) emotions categories based on average rating across participants.

**Table S2.** Final list of movie clips with their description included in the fMRI study.

| Index | Movie | Description of the scene |
|---|---|---|
| 3 | The dead Poets Society | A schoolboy commits suicide |
| 7 | Seven | Policemen find the body of a savagely tortured man |
| 9 | E.T. | E.T. is apparently dying |
| 10 | Trainspotting | Death of a newborn baby |
| 14 | When Harry met Sally | Sally simulates an orgasm in a restaurant |
| 15 | Forrest Gump | Father and son are reunited |
| 16 | Scream | A girl receives threats through the phone |
| 17 | Trainspotting | The "soiled bedsheets" scene |
| 18 | The Professional | The two main characters are separated forever |
| 21 | Dead Man Walking | The main character is put to death by lethal injection |
| 22 | The silence of the lambs | Forensic examination of a dead body |
| 23 | A fish called Wanda | One of the characters (John Cleese) is found naked by the owners of the house |
| 25 | Sleepers | Sexual abuse of children |
| 26 | When a man loves a woman | Reconciliation between two lovers |
| 27 | Saving Private Ryan | Graphic war scene: fighting on the beaches |
| 28 | The Shining | The character played by Jack Nicholson pursues his wife with an axe |
| 30 | In the name of the father | Violent police interrogation leading to forged confessions |
| 31 | Indiana Jones | Indiana Jones (Harrison Ford) escapes from a catacomb full of rats |
| 32 | Copycat | One of the characters gets caught by a murderer in a toilet |
| 33 | Ghost | The "pottery" scene |
| 36 | City of angels | Maggie (Meg Ryan) dies in Seth's (Nicolas Cage) arms |
| 39 | The Piano | One of the characters gets her hand cut off |
| 43 | A perfect World | Butch (Kevin Costner) is gunned down, at the end of the movie |
| 45 | Blue | A man clears out the drawers of his desk; a woman arrives walking in an alley. She greets another woman and continues walking. |
| 46 | Child's Play II (Chucky II) | Chucky beats Andy's teacher with a ruler |
| 48 | Life is beautiful | In a concentration camp, a father "fakes" a translation of what an officer says in order to prevent his son to be frightened |
| 49 | Blue | A woman walks down a street market |
| 50 | Misery | Annie (Kathy Bates) breaks Paul's legs (James Caan) |
| 51 | Leaving Las Vegas | The main character is raped and beaten by three drunk men |
| 52 | Dangerous mind | Students in a school class are told that one of their classmates has died |
| 53 | Underground | A man's attempts to have sex with a prostitute are interrupted by the bombing of the city during WWII |
| 55 | The Blair Witch Project | Final scene in which the characters are apparently killed |
| 56 | Benny and Joone | Benny (Johnny Depp) plays the fool in a coffee shop |
| 57 | Hellraiser | On the floor, the size of two stains are growing, and progressively transforming into a monster with a human-like skeleton |
| 58 | The lover | Marguerite (Jane March) gets into a car, and the car starts to ride. She is dropped of on an animated street. She knocks on a door, and a Chinese man opens and lets her in. |
| 59 | The dead Poets Society | By the end of the movie, all the students climb on their desks to manifest their solidarity with Mr. Keating (Robin William), who has just been fired |
| 61 | There is something about Mary | Ben Stiller fights with a dog |
| 62 | Philadelphia | Andrew (Tom Hanks) and Joe (Denzel Washington) listen to an opera aria on the stereo. Ted describes to Joe the pain and passion felt by the opera character |
| 64 | Blue | A person passes a piece of aluminium foil through the window of a car |

| Category | |
|---|---|
| 1 fear | - |
| 2 disgust | |
| 3 pain | |
| 4 sadness | |
| 5 anger | |
| 6 laughter | + |
| 7 tenderness | |
| 8 neutral | |
| 9 confusion | |

| All videos | | Duration |
|---|---|---|
| 16 | 1 | 409070 |
| 18 | 1 | 171042 |
| 28 | 1 | 266051 |
| 46 | 1 | 67077 |
| 55 | 1 | 246098 |
| 7 | 2 | 109015 |
| 22 | 2 | 218021 |
| 31 | 2 | 118022 |
| 57 | 2 | 96000 |
| 10 | 3 | 98013 |
| 21 | 3 | 416049 |
| 27 | 3 | 335066 |
| 50 | 3 | 221080 |
| 9 | 4 | 286045 |
| 36 | 4 | 268056 |
| 43 | 4 | 278088 |
| 52 | 4 | 135060 |
| 62 | 4 | 342016 |
| 25 | 5 | 146066 |
| 30 | 5 | 210033 |
| 32 | 5 | 151065 |
| 39 | 5 | 43012 |
| 51 | 5 | 156038 |
| 14 | 6 | 171091 |
| 23 | 6 | 173025 |
| 56 | 6 | 125095 |
| 61 | 6 | 219050 |
| 15 | 7 | 128095 |
| 26 | 7 | 100027 |
| 33 | 7 | 216036 |
| 48 | 7 | 228016 |
| 59 | 7 | 159062 |
| 45 | 8 | 40020 |
| 49 | 8 | 25032 |
| 58 | 8 | 43000 |
| 64 | 8 | 16024 |
| 3 | 9 | 253087 |
| 17 | 9 | 64056 |
| 53 | 9 | 58024 |
| Total duration: | | 113.413717 |

| Run1 | Category | Duration | |
|---|---|---|---|
| 16 | 1 | 409070 | |
| 10 | 3 | 98013 | |
| 36 | 4 | 268056 | |
| 25 | 5 | 146066 | |
| 14 | 6 | 171091 | + |
| 15 | 7 | 128095 | + |
| 49 | 8 | 25032 | + |
| 53 | 9 | 58024 | + |
| | | 21.72 | min |

| Run2 | Category | Duration | |
|---|---|---|---|
| 18 | 1 | 171042 | |
| 7 | 2 | 109015 | |
| 21 | 3 | 416049 | |
| 52 | 4 | 135060 | |
| 30 | 5 | 210033 | |
| 23 | 6 | 173025 | + |
| 26 | 7 | 100027 | + |
| 58 | 8 | 43000 | + |
| | | 22.62 | min |

| Run3 | Category | Duration | |
|---|---|---|---|
| 28 | 1 | 266051 | |
| 57 | 2 | 96000 | |
| 27 | 3 | 335066 | |
| 43 | 4 | 278088 | |
| 32 | 5 | 151065 | |
| 56 | 6 | 125095 | + |
| 59 | 7 | 159062 | + |
| 64 | 8 | 16024 | + |
| | | 23.77 | min |

| Run4 | Category | Duration | |
|---|---|---|---|
| 46 | 1 | 67077 | |
| 22 | 2 | 218021 | |
| 50 | 3 | 221080 | |
| 9 | 4 | 286045 | |
| 39 | 5 | 43012 | |
| 61 | 6 | 219050 | + |
| 48 | 7 | 228016 | + |
| 17 | 9 | 64056 | + |
| | | 22.44 | min |

| Run5 | Category | Duration | |
|---|---|---|---|
| 55 | 1 | 246098 | |
| 31 | 2 | 118022 | |
| 62 | 4 | 342016 | |
| 51 | 5 | 156038 | |
| 33 | 7 | 216036 | + |
| 45 | 8 | 40020 | + |
| 3 | 9 | 253087 | + |
| | | 22.86 | min |

**All the clips**

| | Amount | Duration |
|---|---|---|
| - | 23 | 79.77 |
| + | 16 | 33.65 |

**Best randomized sequences so far**

| | | | | | | | | |
|---|---|---|---|---|---|---|---|---|
| Run1 | 49 | 36 | 10 | 16 | 53 | 15 | 25 | 14 |
| | 8 | 4 | 3 | 1 | 9 | 7 | 5 | 6 |
| Run2 | 26 | 18 | 30 | 52 | 23 | 58 | 7 | 21 |
| | 7 | 1 | 5 | 4 | 6 | 8 | 2 | 3 |
| Run3 | 32 | 27 | 64 | 28 | 56 | 57 | 43 | 59 |
| | 5 | 3 | 8 | 1 | 6 | 2 | 4 | 7 |
| Run4 | 22 | 48 | 61 | 9 | 39 | 46 | 50 | 17 |
| | 2 | 7 | 6 | 4 | 5 | 1 | 3 | 9 |
| Run5 | 62 | 3 | 51 | 33 | 31 | 55 | 45 | |
| | 4 | 9 | 5 | 7 | 2 | 1 | 8 | |

* neutral and confusion are assigned to positive just because they are not negative

**Figure S2.** Assignment of the movie clips into the 5 running sessions (runs) based on emotional content.

### 1.2. Emotional ratings

The same 39 movie clips divided into 3-10 seconds long parts, were rated later by 16 Finnish-speaking female raters on scale 0-4 in order to evaluate the intensity of 63 perceived and experienced emotions (in Finnish). The ratings were done with online own developed tool (**Figure S3**). The participants rated the movies in a random order, while the order of the scenes was retained. The participants were encouraged to rate all scenes of a movie one after another to enable the rating of slowly developing emotions (if participants were idle for a long period, a popup appeared to suggest that they restart the movie from the first scene). However, the participants were also encouraged to have breaks between movies to avoid fatigue during the long rating process. The raters were also compensated for their effort.

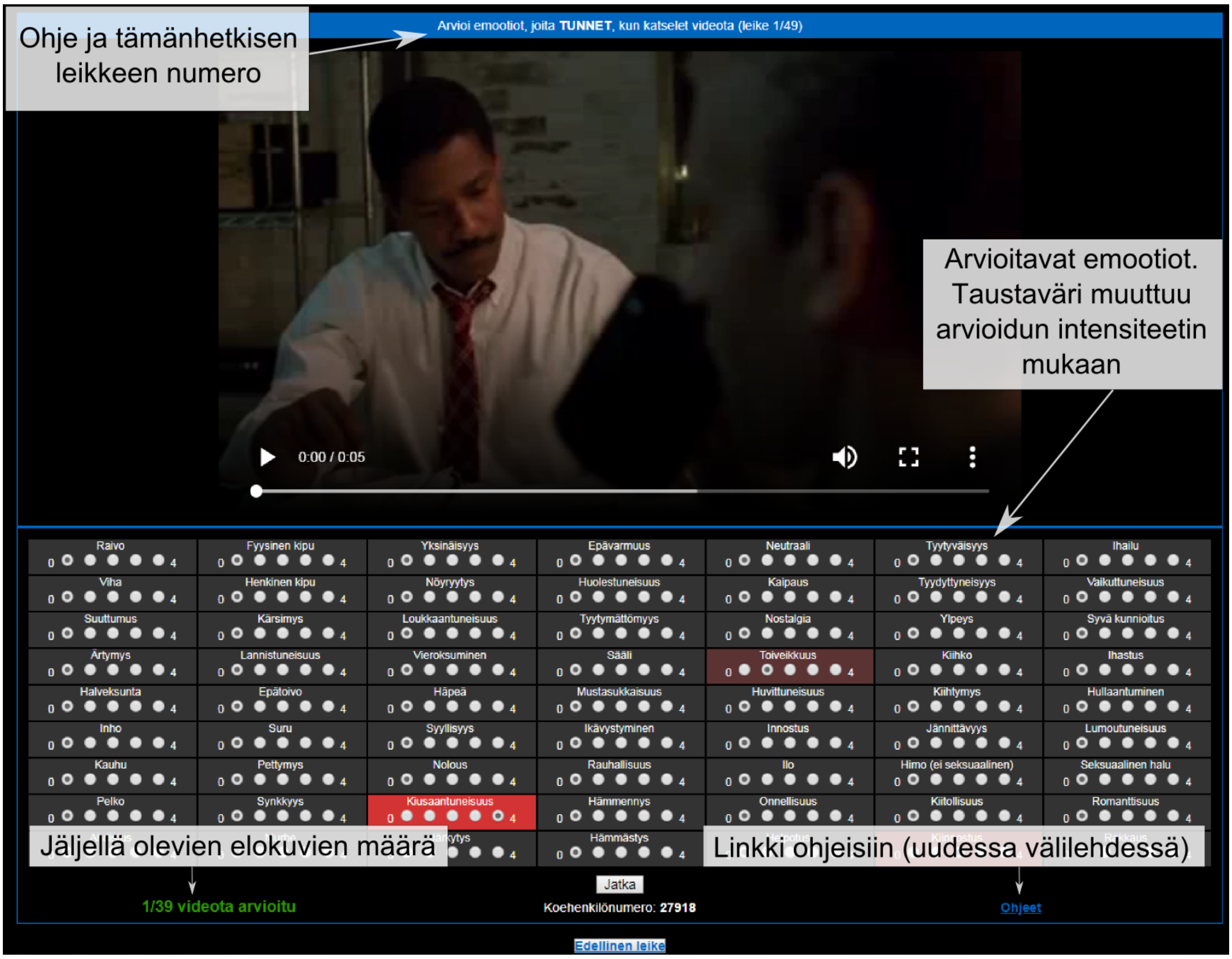


**Figure S3.** The example of rating tool as a tutorial for the subject (In Finnish).

#### 1.3. Stimulus presentation and physiological measures

The auditory track of the movies was delivered through Sensimetrics S14 insert earphones (Sensimetrics Corporation, Malden, Massachusetts, USA). Sound was adjusted for each subject to be loud enough to be heard over the scanner noise. The visual stimuli were back-projected on a semitransparent screen using a Panasonic PT-DZ110XEJ data projector (Panasonic, Osaka, Japan) and from there viewed by the participant via a mirror fixed to the head coil. Stimulus presentation was controlled with Presentation software (Neurobehavioral Systems Inc., Albany, CA, USA). Heart rate and respiration data was collected successfully from XX subjects with BIOPAC MP150 Data Acquisition System (BIOPAC System, Inc., USA). Heart rate was measured using BIOPAC TSD200 pulse plethysmogram tranducer, which records the blood volume pulse waveform optically. The pulse transducer was placed on the palmar surface of the participant's left index finger. Respiratory movements were measured using BIOPAC TSD201 respiratory-effort transducer attached to an elastic respiratory belt, which was placed around each participant's chest to measure changes in thoracic expansion and contraction during breathing. Both signals were sampled simultaneously at 1 kHz using RSP100C and PPG100C amplifiers for respiration and heart rate, respectively, and BIOPAC AcqKnowledge software (version 4.1.1).

#### 1.4. MRI data acquisition and processing

MRI data were collected using Siemens Skyra 3T MRI with 30-channel head coil at the Advanced Magnetic Imaging Centre, Aalto Neuroimaging, Aalto University. High-resolution anatomical images ( 1 x 1 x 1 mm voxel size) were acquired using T1-weighted MP-RAGE sequence with 2530-ms TR. Functional MRI was obtained with whole-head T2*-weighted echo-planar imaging pulse sequence (TR=2400 ms, TE=24 ms, flip angle=70°, and 3 x 3 x 3 mm resolution; fat-suppression with bipolar water excitation radio frequency pulse.

fMRI data were preprocessed using FMRIPREP (v016216) (Gorgolewski et al., 2011; Esteban et al., 2019). Each T1-weighted volume was corrected for intensity non-uniformity using N4BiasFieldCorrection (v2.1.0) and skull-stripped using ANTs (v2.1.0; the OASIS template). Brain surfaces were reconstructed using recon-all from FreeSurfer (v6.0.1) which also produced volume estimates of cortical (34 for each hemisphere) and subcortical (63) brain structures for each subject. The brain mask was refined with a custom variation of the method to reconcile ANTs-derived and FreeSurfer-derived segmentations of the cortical gray matter of Mindboggle (v002438). Spatial normalization to the ICBM 152 Nonlinear Asymmetrical template v2009c was performed through nonlinear registration with the antsRegistration tool of ANTs, using brain-extracted versions of both

T1w volume and template. Brain tissue segmentation of cerebrospinal fluid, white-matter, and gray-matter was performed on the brain-extracted T1 using FSL Fast (v5.0.9). Functional data was slice time corrected using 3dTshift from AFNI (v16.2.07), motion corrected using MCFLIRT (v5.0.9), followed by co-registration to anatomical scans using boundary-based registration with 9 degrees of freedom in FreeSurfer. Motion correction transformations, BOLD-to-T1w transformation and T1w-to-template (MNI) warp were concatenated and applied in a single step using ANTs with Lanczos interpolation.

Physiological noise regressors were extracted applying CompCor (Behzadi et al., 2007). Principal components were estimated for the two CompCor variants: temporal (tCompCor) and anatomical (aCompCor). A mask to exclude signal with cortical origin was obtained by eroding the brain mask, ensuring it only contained subcortical structures. Six tCompCor components were then calculated including only the top 5% variable voxels within that subcortical mask. For aCompCor, six components were calculated within the intersection of the subcortical mask and the union of CSF and WM masks calculated in T1w space, after their projection to the native space of each functional run. Framewise displacement was calculated for each functional run using the implementation in Nipype.

After FMRIPREP, normalized data was denoised using Bramila tools (https://version.aalto.fi/gitlab/BML/bramila) including temporal detrending (Savitzky-Golay filter with 240s window; Press and Teukolsky, 1990), nuisance regression, temporal filtering (high-pas 0.0067Hz) and spatial smoothing (6mm Gaussian kernel). The nuisance regression on task-related data included 6 motion estimators (3 translations + 3 rotations), CSF and WM signals.

#### 1.5. Data preparation and analyses

Firstly, we checked that it is possible to differentiate subjects based on high and low scoring in PP and SP (**Figure S4** and **S5)**. Secondly, the ratings of all 63 experienced and perceived emotions were inspected visually for consistency as a) variation in ratings across all movies (**Figure S6** and **S7**) and b) amount of non-zero ratings per emotion (**Figure S8** and **S9**). Then the rated perceived and felt emotions were evaluated by inter-raters agreement (IRR; **Figure S10**), based on which we chose 6 basic perceived emotions *(fear, sadness, happiness, anger, surprise, disgust)* and neutral stimulus as reliably rated and relevant for psychopathy research.

First, we concatenated preprocessed and smoothed fMRI BOLD activity over the length of all movie stimuli for each of 50 subjects. We then generated the Intersubject correlation (ISC) similarity matrices using Bramila tools (https://version.aalto.fi/gitlab/BML/bramila). The ISC matrices showed whole-brain voxel-wise correlations in brain activations across subjects. Next, we performed ISC-based two-sample t-test using Bramila with 25-75 percentile group split and element-wise permutations. The local maxima were estimated over found brain clusters (min. size 105 voxels) in regions from Brainnetome and Cerebellar Connectivity atlases with total 334 structures. The results with low and high PP/SP mean regional activity over the maxima were presented on cluster corrected brain maps with $p<0.0001$.

Second, the preprocessed and smoothed fMRI BOLD activity data was overlapped (interpolated) over higher than zero (non-zero) rated time-points of movies for each of basic perceived emotions and neutral stimuli. The fMRI BOLD data was then concatenated for each emotion and subject, the ISC matrices were obtained and ISC-based t-test was performed in the same manner as previously with the cluster corrected brain maps with $p<0.0001$ as outcome.

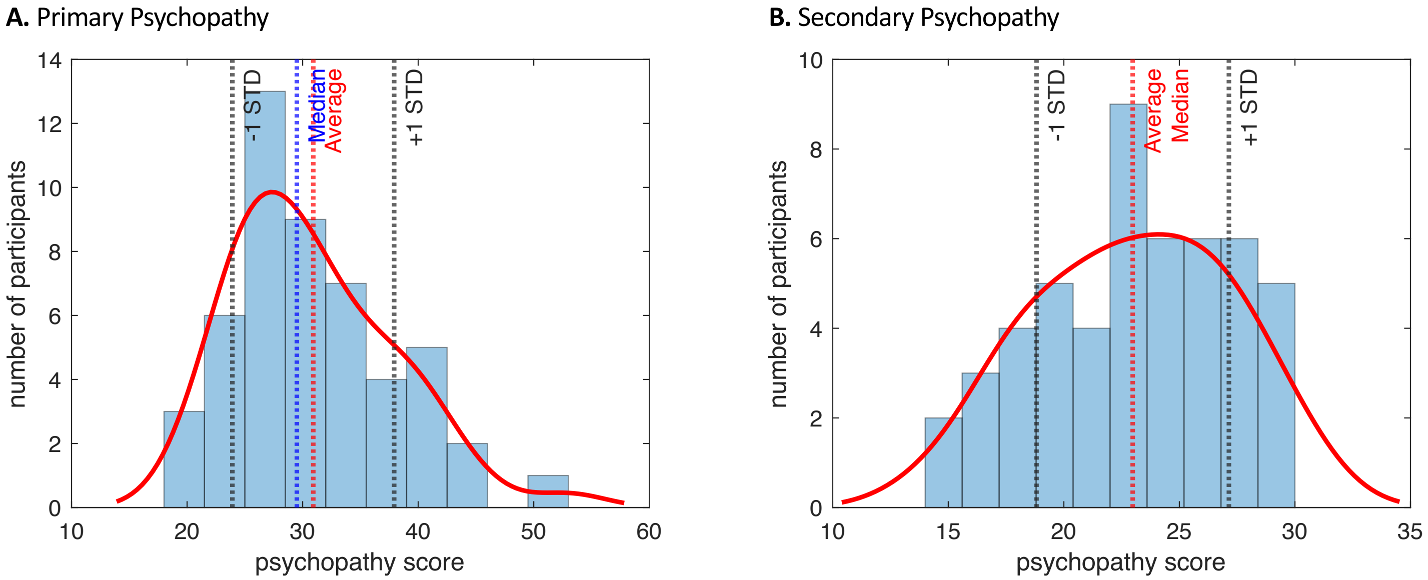


**Figure S4.** Distributions of participants' scores on primary psychopathy scale (A) and secondary psychopathy scale (B) from Levenson Self-Report Psychopathy Scale. The plots show median and average psychopathy scores in each scale along with 1 standard deviations. The red curve is a kernel function fitted onto the data. The scores obtained from primary psychopathy scale present right skewed distribution (A) and scores from secondary psychopathy scale normal distribution (B).

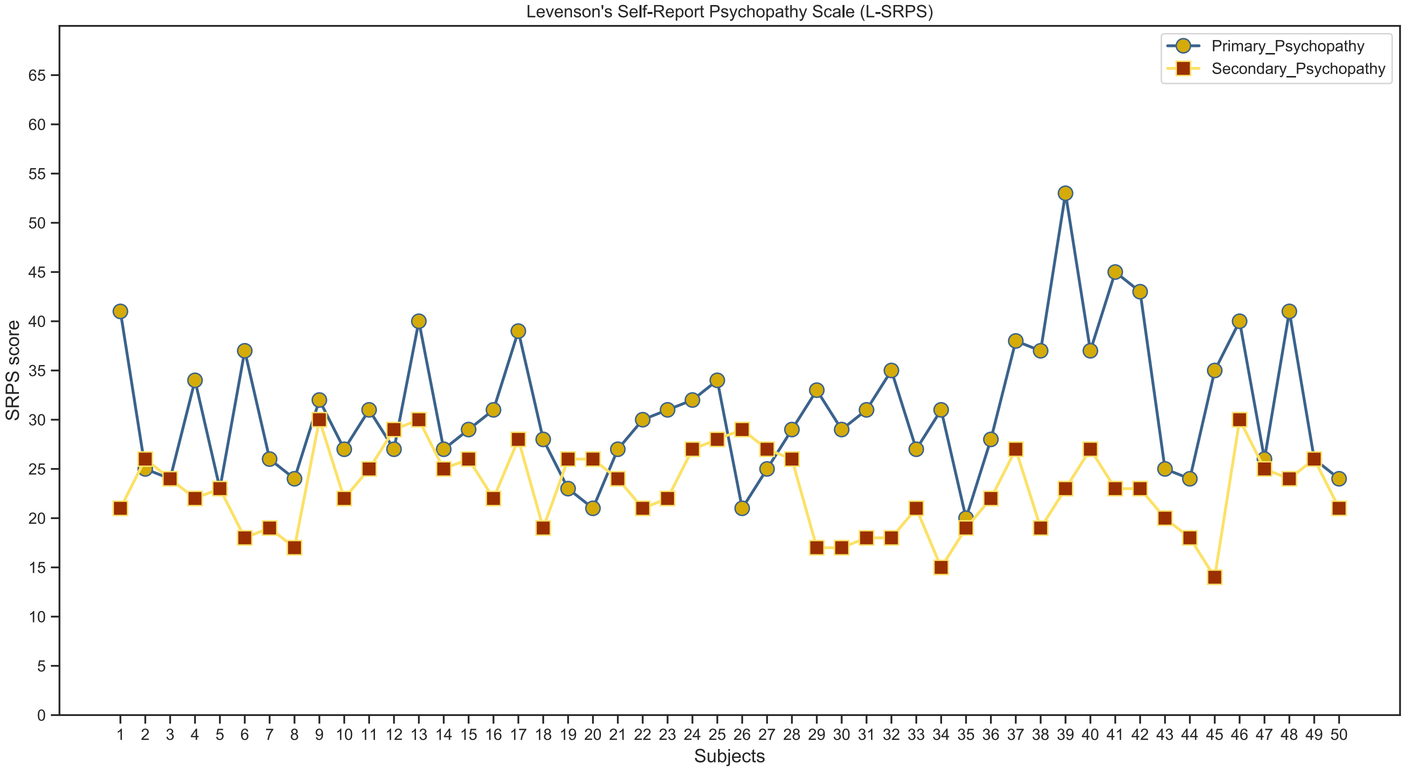


**Figure S5.** Psychopathy score among the subjects. Graph presents the variation in individual scores of 50 participants in primary and secondary psychopathy based on Levenson's Self-Report Psychopathy Scale (LSRPS). The higher the score, the more psychopathy traits the person owns.

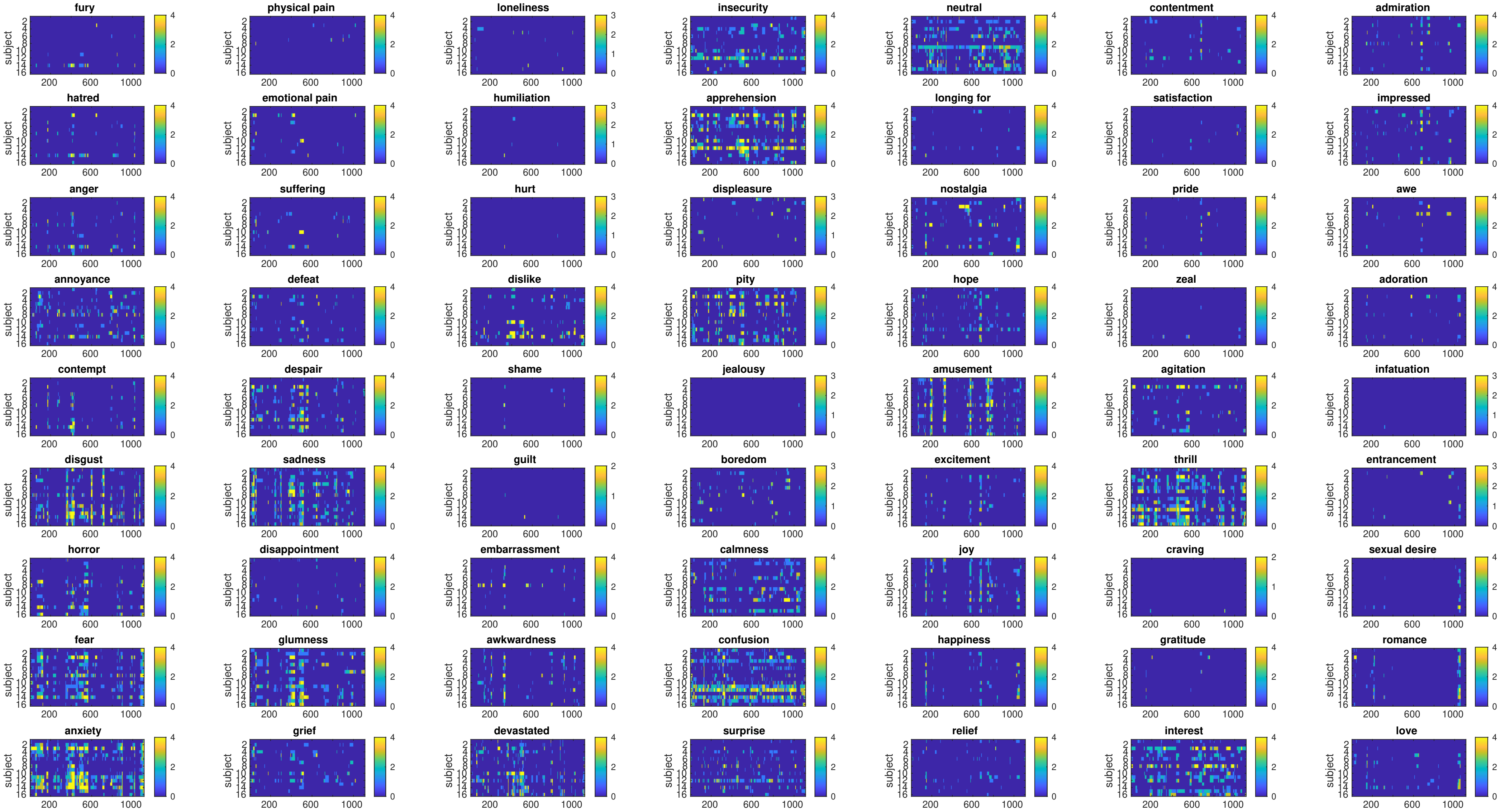


**Figure S6.** Ratings of experienced emotions across subjects. The averaged scoring (0-4) of each movie's timepoints across the raters.

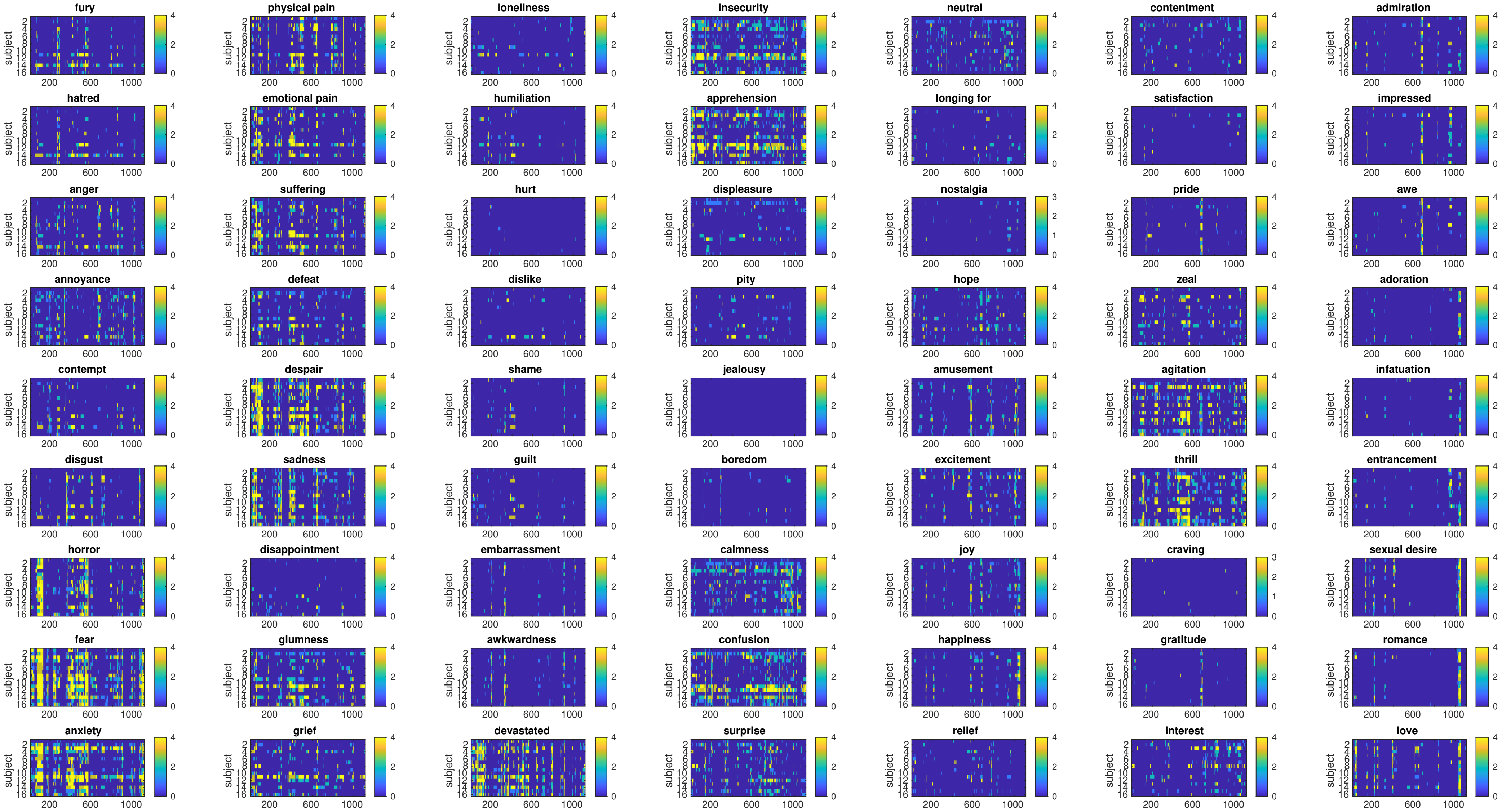


**Figure S7.** Ratings of perceived emotions across subjects. The averaged scoring (0-4) of each movie's timepoints across the raters.

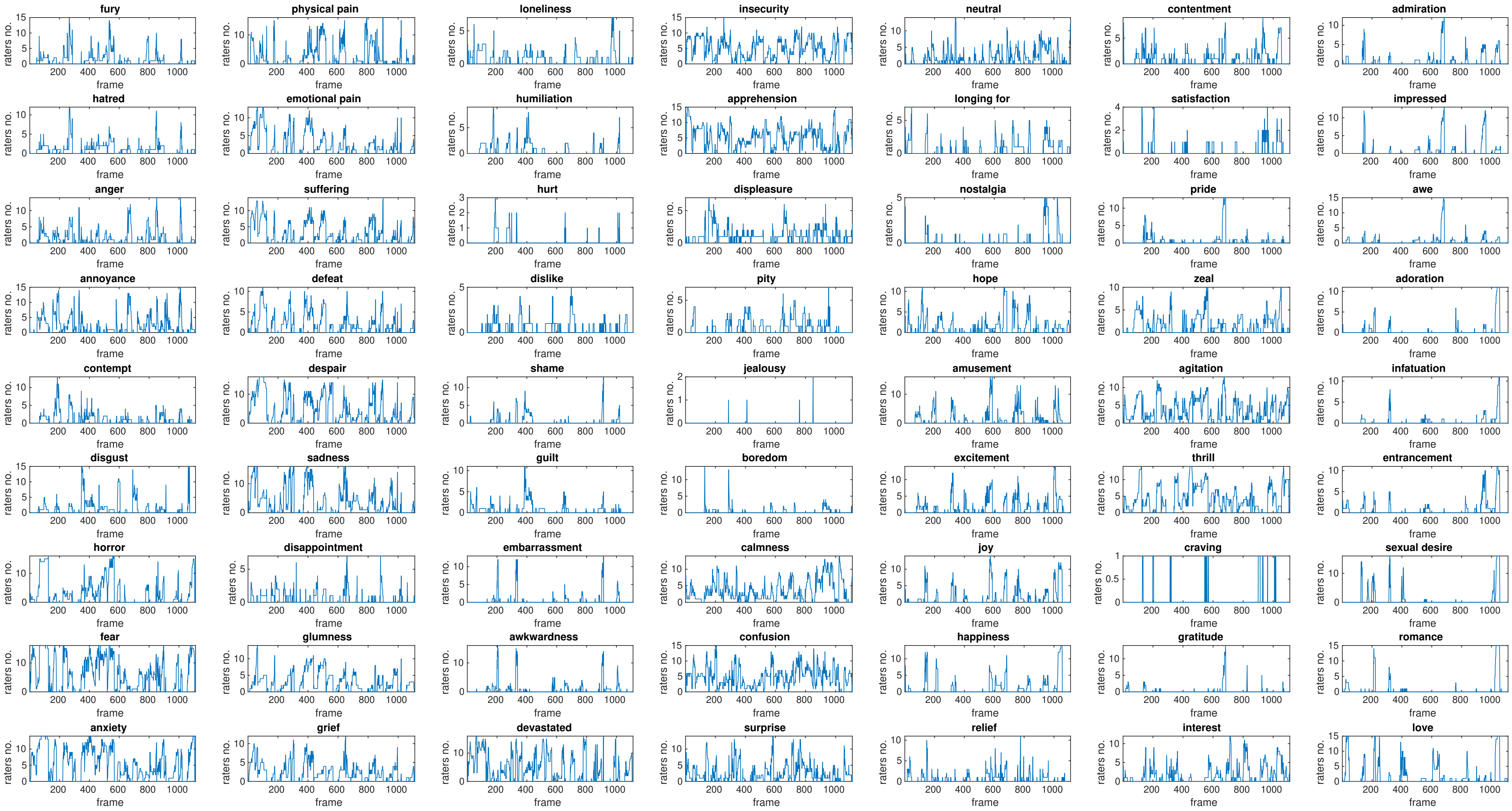


**Figure S8.** Number of raters with non-zero ratings in experienced emotions. Averaged non-zero ratings of each movie timepoint per emotion across subjects.

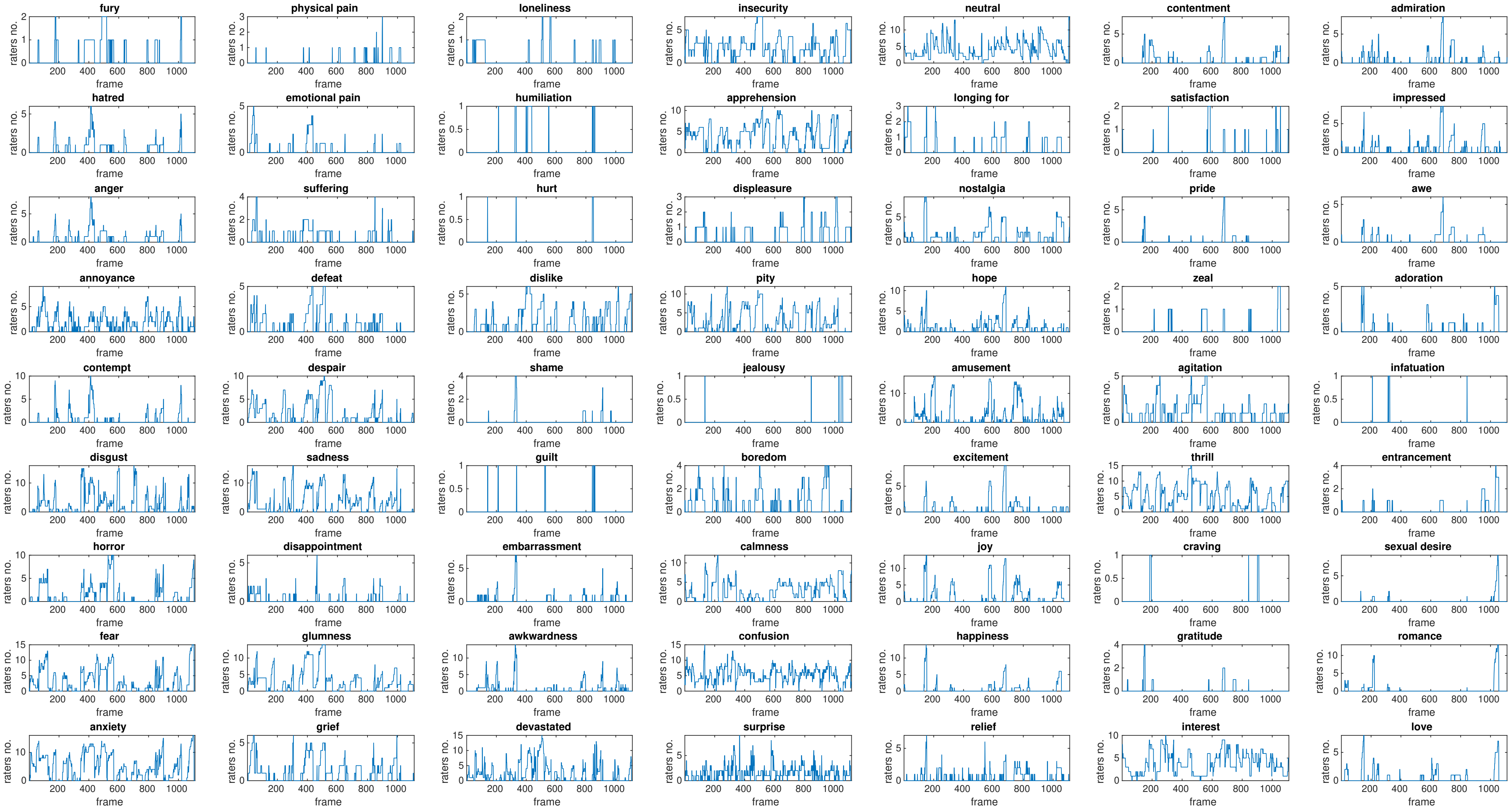


**Figure S9.** Number of raters with non-zero ratings in perceived emotions. Averaged non-zero ratings of each movie timepoint per emotion across subjects

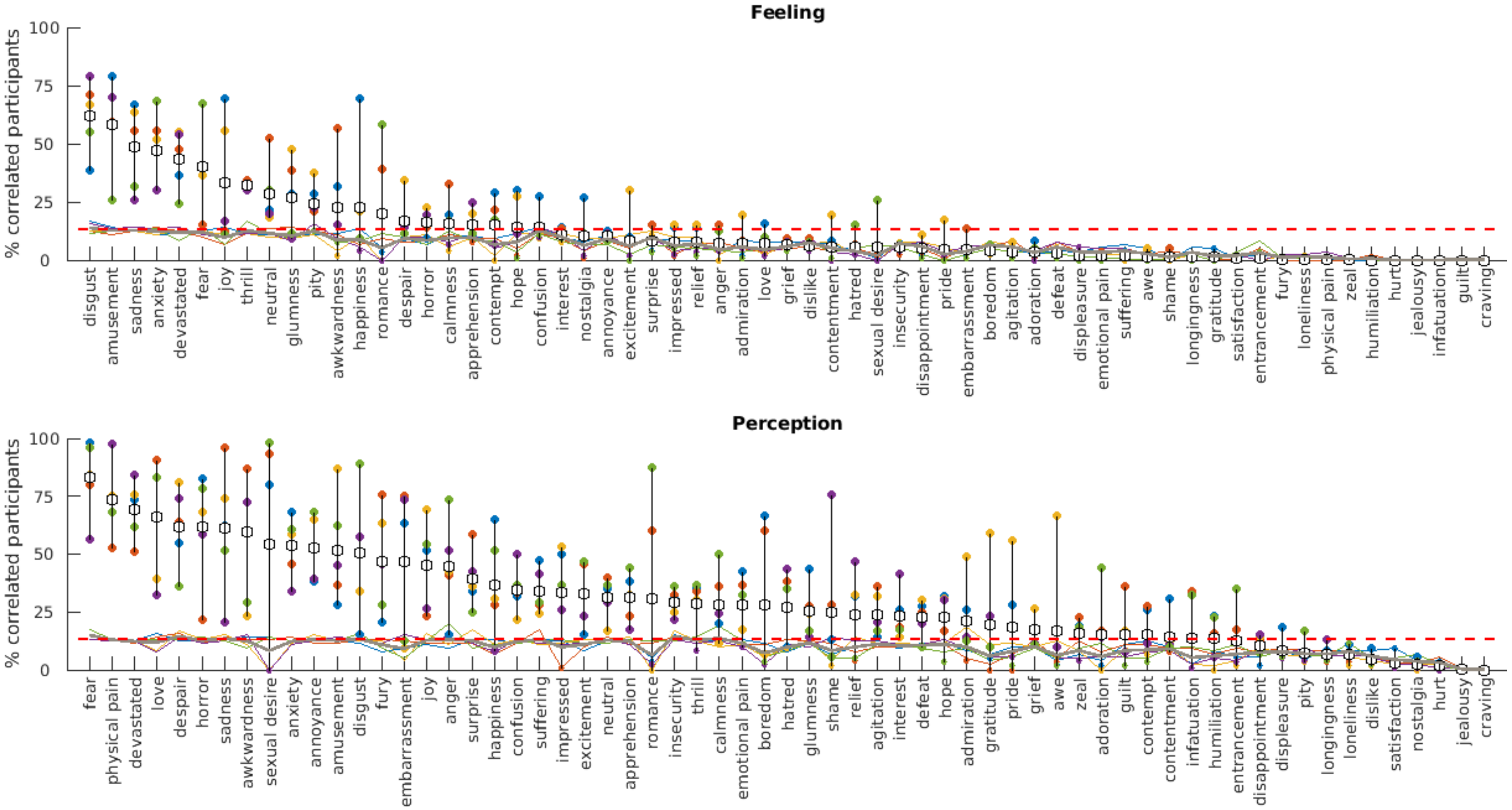


**Figure S10.** Inter-raters reliability (IRR) of emotional content in the movies. Inter-raters agreement between subjects scoring experienced and perceived emotions in the movie clips. The plots showing agreement between raters as a percentage of significantly correlated pairs (Spearman) sorted by highest values. Color lines show individual permuted correlation thresholds for each emotion and for each of 5 runs of movies; thick grey line is mean threshold for each emotion over 5 runs and dashed red line is Bonferroni threshold, pairs corrected for multiple comparisons. Color dots show percentage of correlated pairs for each of 5 runs of movies and white dots are means over 5 runs. In order to take emotion as reliable it must pass the Bonferroni threshold.
